# Interpolating and Extrapolating Node Counts in Colored Compacted de Bruijn Graphs for Pangenome Diversity

**DOI:** 10.64898/2026.03.16.711983

**Authors:** Luca Parmigiani, Pierre Peterlongo

**Affiliations:** Faculty of Technology and Center for Biotechnology (CeBiTec), Bielefeld University, Bielefeld, Germany; University Rennes, Inria, CNRS, IRISA - UMR 6074, Rennes, France

**Keywords:** Pangenomics, pangenome graphs, colored compacted de Bruijn graphs, Hill numbers

## Abstract

A pangenome is a collection of taxonomically related genomes, often from the same species, serving as a representation of their genomic diversity. The study of pangenomes, or pangenomics, aims to quantify and compare this diversity, which has significant relevance in fields such as medicine and biology. Originally conceptualized as sets of genes, pangenomes are now commonly represented as pangenome graphs. These graphs consist of nodes representing genomic sequences and edges connecting consecutive sequences within a genome. Among possible pangenome graphs, a common option is the compacted de Bruijn graph. In our work, we focus on the colored compacted de Bruijn graph, where each node is associated with a set of colors that indicate the genomes traversing it.

In response to the evolution of pangenome representation, we introduce a novel method for comparing pangenomes by their node counts, addressing two main challenges: the variability in node counts arising from graphs constructed with different numbers of genomes, and the large influence of rare genomic sequences. We propose an approach for interpolating and extrapolating node counts in colored compacted de Bruijn graphs, adjusting for the number of genomes. To tackle the influence of rare genomic sequences, we apply Hill numbers, a well-established diversity index previously utilized in ecology and metagenomics for similar purposes, to proportionally weight both rare and common nodes according to the frequency of genomes traversing them.

## 1 Introduction

A pangenome is the collection of all genomes from the same taxonomic ranking, like species or genus. Because the number of organisms on Earth is finite, the number of possible genomes is also finite, regardless of their vastness (Mushegian 2020; Landenmark et al. 2015). Thus, the question of how faithfully our sampled pangenome represents the real pangenome boils down to its diversity. If our sequenced genomes exhibit little variation, a relatively small number of genomes might suffice. Conversely, significant variation would necessitate sequencing a larger number of genomes. Current data suggest the latter is true for most pangenomes. In fact, most genetic material, such as genes, is rare, leading to incomplete pangenomes. This was first realized by Tettelin et al. (2005), who predicted that the number of genes would grow indefinitely with the number of genomes sequenced for some bacterial species. Although this value cannot reach infinity in practice (Haegeman and Weitz 2012), it serves as a useful theoretical model, analogous to the infinitely many alleles model (Ewens 2004), reflecting the considerable variation in genetic content observed across genomes of the same species (Tettelin et al. 2005).

Nowadays, pangenomes are commonly represented as pangenome graphs (Eizenga et al. 2020), where nodes are labeled with genetic sequences, and edges connect consecutive sequences within genomes. Modeling pangenomes as graphs offers two main advantages: first, they compress sequences shared among genomes, and second, they allow the identification of variations between genomes. This immediately raises questions about the extent of variation within a pangenome and the degree of shared genetic material among genomes. Although many methods exist for graph comparison, it is essential to recognize that a pangenome graph is a subset of a much larger graph representing the pangenome of all genomes from a species or genus.

In this context, graph comparison suffers from two well-known problems in ecology: a *sampling problem* and an *abundance problem*. The sampling problem arises because pangenome graphs are often constructed from different numbers of genomes, leading to differences in the number of nodes and edges, making direct comparisons difficult. The abundance problem refers to the strong influence of rare genomic sequences, since many sequences occur in only a few genomes and thus inflate the apparent diversity. A common way to address the sampling problem is to standardize diversity estimates to a fixed number of genomes using interpolation and extrapolation. To reduce the influence of rare sequences, diversity measures that account for genomic element frequencies can be used.

The simplest diversity measure is *richness*, defined as the total number of distinct genomic elements observed. In pangenomes, richness usually corresponds to the number of unique genetic elements such as genes or *k*-mers. While species richness has received considerable attention (Hogg et al. 2007; Sheikhizadeh et al. 2016; Page et al. 2015; Parmigiani et al. 2024), and was a focus since the first pangenome paper (Tettelin et al. 2005), the use of Hill numbers to compare pangenomes has remained underutilized in pangenomics. This is not entirely surprising, as a similar situation occurred in ecology, where Hill numbers and species richness (or rarefaction) developed independently and were only later unified by Chao et al. (2014).

In this work, we extend this type of diversity analysis to colored compacted de Bruijn graphs, a type of graph that can be used as a pangenome graph. We present a novel analytical formula for the interpolation and a nonparametric estimator for the extrapolation of the number of nodes, designed to be used with Hill numbers. Our results demonstrate that the interpolation method is faster and more robust than building the pangenome graph for multiple permutations of the order of genomes. We also evaluate the effectiveness of the nonparametric estimator. Finally, we compared 12 bacterial pangenomes represented as colored compacted de Bruijn graphs. The code was implemented in the tool Pangrowth, which is available at https://github.com/gi-bielefeld/pangrowth.

## 2 Preliminaries

### Strings

Let *s* ∈ Σ^∗^ be a string over a totally ordered alphabet Σ, and let *s*[*i* : *j*] denote the substring of *s* that starts at index *i* and ends at index *j*. The element at position *i* is denoted as *s*[*i*]. The lexicographic order ≤_lex_ follows standard dictionary ordering: given two strings *s* and *t*, we say that *s* is lexicographically smaller or equal than *t* if either *s* is a prefix of *t* or there exists an index *i* such that the first *i* − 1 characters of both strings are identical, but the *i*th character of *s* is smaller than the *i*th character of *t*. Let $ ∈*/* Σ be a sentinel character that is considered smaller than any *c* ∈ Σ. The sentinel is often appended at the end or at the start of the string (e.g., *s*$ or $*s*). We define a *genome G* as a set of strings. A *pangenome G* is a set of *N* genomes:*G* = {*G*_1_, …, *G*_*N*_}.

Let *k* be a positive integer, a *k-mer* is any substring of *s* of length *k*, denoted *s*[*i* : *i* + *k* − 1]. The *spectrum* of a string *s*, denoted *S*_*k*_(*s*), is the set of all distinct *k*-mers present in *s*. The set *S*_*k*_(*G*) represents all distinct *k*-mers occurring in at least one string of a genome in *G*. In DNA, a sequence and its reverse complement are considered equivalent. For a string *s* = *s*_1_*s*_2_ … *s*_*n*_, the reverse complement is obtained by reversing the sequence and replacing each nucleotide with its paired base (*A* ↔ *T*, *C* ↔ *G*) *s*^*rc*^ = *c*(*s*_*n*_) *c*(*s*_*n*−1_) … *c*(*s*_1_), where *c* denotes the base-pair mapping. To ensure a unique representation of each sequence regardless of strand orientation, we define a *canonical k*-mer 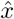 as the lexicographically smaller of *x* and its reverse complement 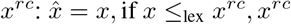, otherwise. The set of canonical *k*-mers of a string *s* is referred to as the *canonical spectrum*.

### Colored compacted de Bruijn graph

A *graph* is a pair of sets (*V, E*), where *V* is a set of *nodes* and *E* is a set of pairs of nodes, called *edges*. A graph is called *directed* if the order of the edge matters, i.e., (*v, u*) ≠ (*u, v*). Given a pangenome *G*, the *de Bruijn graph dBG*(*G*) of order *k* is the directed graph with a node for each *k*-mer present in a genome in *G* and an edge (*u, v*) exists if *u*[2 : *k*] = *v*[1 : *k* − 1], which we sometimes refer to as “*u* overlaps with *v*”.

A *compacted de Bruijn graph cdBG*(*G*) is a graph obtained from a de Bruijn graph where each non-branching path, that is, a maximal path whose internal nodes have in degree and out degree equal to one, is represented as a single node called *unitig*. The process of merging two *k*-mers in a single one is called *compaction*. A *colored compacted de Bruijn graph ccdBG*(*G*) is a compacted de Bruijn graph where each unitig is associated with a set of colors, each representing a genome that contains that unitig. Some models allow a genome to be associated with a unitig even if it contains only a subset of the *k*-mers within it. Here, we assume that unitigs are uni-chromatic, meaning that each unitig is composed entirely of *k*-mers that share the same color set. Due to the double-stranded nature of DNA, a de Bruijn graph can be built using canonical *k*-mers. In this case there is an edge (*u, v*) if *u*[2 : *k*] = *v*[1 : *k* − 1] or *u*[2 : *k*] = *v*^*rc*^[1 : *k* − 1], where *v*^rc^ denotes the reverse complement of *v*. The compaction remains unchanged. We refer to these graphs as canonical de Bruijn graphs (or canonical cdBG/ccdBG).

### Genome as items

We now abstract a genome as a set of *items*. Items are genomic features represented by their presence or absence across genomes, and can occur in one or more genomes. In classical pangenomic analyses, these items are typically the genes contained within a genome. However, many alternatives are possible, for instance *k*-mers (Parmigiani et al. 2024), minimizers (Ekim et al. 2021), or open reading frames (ORFs) (Vernikos et al. 2015).

Let *t* denote the item type. Given a genome *G* ∈ *G, G*^*t*^ is the set of items of type *t* extracted from *G*. Each *G*^*t*^ represents a subset of the universal set of items *I*^*t*^ present in a species or other taxonomic rank. The set *I*^*t*^ is defined as all items observed in the pangenome plus any unseen items that may appear in genomes not yet included. This universe is not necessarily equal to the full combinatorial space of possible items. For example, for DNA *k*-mers, the full combinatorial space has size 4^*k*^, but not all 4^*k*^ strings need to occur in genomes from the considered species. We refer to the size of *I*^*t*^ as *I*^*t*^. The item representation of a pangenome is then 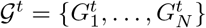. For example, when items are *k*-mers, the universe *I*^*k*-mer^ contains all *k*-mers that appear in the considered species. When the type of item is not relevant, we simply write *I* and *I*.

We define a *pan-matrix M*, with *N* columns, one for each genome, and *I* rows, one for each possible item. Each entry *M*_*xi*_ equals 1 if item *x* is present in genome 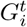, and 0 otherwise. The *incidence Y*_*x*_ of an item *x* is defined as the number of genomes in which the item appears 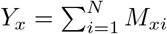. This value indicates how common an item is in the pangenome.

To understand the overall distribution of item incidences, we define the item *frequency h*(*i*) as the number of items appearing in *exactly i* genomes: *h*(*i*) = |{*x* ∈ *I* | *Y*_*x*_ = *i*}|. The value of *h*(1) corresponds to the number of unique items, *h*(2) the number of items that appear in two genomes, and so on. The value *h*(0) represents the number of items that were not seen in any of the *N* genomes but are present in some genome not yet sequenced. This number is usually unknown, but as we will see, it is important for extrapolating the total number of items *I*. For a sample of *m* genomes, let *H*_*m*_(*i*) be the random number of items observed in exactly *i* of the *m* genomes, and let *h*_*m*_(*i*) = E[*H*_*m*_(*i*)] denote its expectation.

We model a pangenome *G* from a probabilistic perspective in the following way. When the presence of each item *x* in a genome follows a Bernoulli trial with a uniform probability *π*_*x*_ for each genome *P* (*M*_*xi*_ = 1) = *π*_*x*_ and *P* (*M*_*xi*_ = 0) = 1 − *π*_*x*_.

Note that ∑_*x*∈*I*_*π*_*x*_ can be greater than one. Let 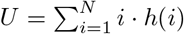 be the total number of items found, counted with their multiplicity. The expected number of items that appear in *N* genomes follows (Chao 1987):

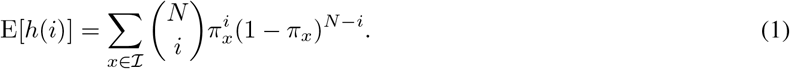

In practice, we are often interested in how the number of items changes with the number of sampled genomes. For example, we may want to estimate the expected number of items for fewer genomes than currently observed, or predict how many additional items would appear if more genomes were sequenced.

These tasks are known as *interpolation* and *extrapolation*. Interpolation estimates the number of items for a subsample drawn from the *N* observed genomes. Extrapolation predicts the number of items for more than *N* genomes, extending the expected accumulation beyond the currently observed genomes.

### Hill numbers

In some species, most genetic material, such as genes, is unique to individual genomes. This leads to pangenomes that are predominantly composed of unique items, making the total number of items *I* an upper bound rather than an accurate reflection of their diversity. It is somewhat counterintuitive that “rare” items, unique to individual genomes, make up the majority of the pangenome.

This problem is also known in other fields, such as ecology, where the objective is to express the diversity of species within an assemblage (a group of co-occurring species). A possible solution is to use Hill numbers (Hill 1973), a family of diversity indices introduced in 1973 and widely used in the field of ecology. Hill numbers can be expressed as a weighted average over a common parameter *q* and they exhibit many intuitive properties, such as the *doubling property*: if two completely distinct pangenomes (i.e., with no shared items) have identical Hill numbers of order *q*, then the Hill number of the combined pangenome is exactly twice the individual value. Moreover, for certain values of *q*, they correspond to well-known diversity measures. Together with interpolation and extrapolation methods, Hill numbers provide a powerful tool to investigate and compare biodiversity.

To calculate Hill numbers, it is essential that the sum of the probabilities equals one. Since ∑_*x*∈*I*_*π*_*x*_ can be greater than one, we normalize each probability to obtain a *relative incidence p*_*x*_ = *π*_*x*_*/*∑_*x*∈*I*_ *π*_*x*_, as proposed by Chao et al. (2014). Hill numbers of order *q* are defined as:

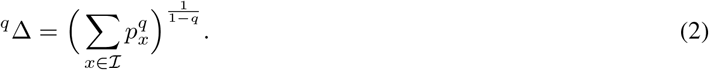

For *q* = 0 the incidence of an item is not affecting the total sum, resulting in the total number of items in a pangenome:

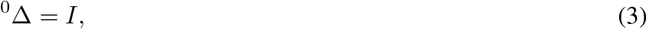

usually referred to as *richness*. For *q* = 1 the formula is undefined, but the limit of *q* → 1 corresponds to the exponential of Shannon’s entropy:

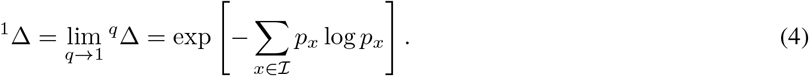

For *q* = 2 it corresponds to the inverse of Simpson’s concentration:

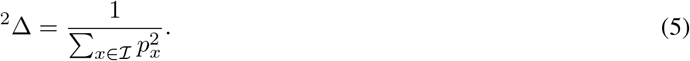

While ^*q*^Δ exists for every value of *q* ∈ ℝ, values of *q >* 2 tend to give too much weight to high probabilities, and they are less used in practice. Let ^*q*^Δ(*m*) represent the expected Hill number of order *q* for *m* genomes. When considering genomes beyond the observed pangenome of size *N*, we write ^*q*^Δ(*N* + *m*^∗^) to denote the estimated Hill number of order *q* for *m*^∗^ *>* 0 additional genomes. As *m* approaches infinity, it converges to ^*q*^Δ(∞) := ^*q*^Δ.

## 3 Uni-mers and infix equivalents

In the previous section, our abstraction allowed for items like *k*-mers and genes. These items are relatively straightforward to analyze because the presence of one *k*-mer (or gene) in one genome has no impact on the presence of another *k*-mer in another genome. However, in this section we shift our focus to unitigs, which are nodes representing non-branching paths in the original de Bruijn graph. When a new sequence is added, unitigs can break into smaller segments or, less frequently, merge into longer ones. This dependency makes the analysis of unitigs different. To make this problem tractable, we show that we can reduce our analysis to *k*-mers and *uni-mers*, a new concept introduced below. For simplicity, throughout most of this work we treat a *k*-mer and its reverse complement as *distinct*. In Subsection 4.1, we show how our method extends to canonical *k*-mers, accounting for the double-stranded nature of DNA.

Given a pangenome *G* and a positive constant *k*, we define the *extended spectrum* of *G* as ℰ_*k*+1_(*G*) = {*xy*[*k*] | *x, y* ∈ *S*_*k*_(*G*) ∧ *x*[2 : *k*] = *y*[1 : *k* − 1] ∧ *x* ≠ *y*} representing the (*k* + 1)-mers that are formed by concatenating a pair of distinct overlapping *k*-mers *x, y* ∈ *S*_*k*_(*G*). The condition *x* ≠ *y* ensures that we consider only transitions between distinct *k*-mers, as we are interested in potential extensions that may compact into a unitig.

Let *h*^unitig^(*i*) be the *unitig frequency* of the ccdBG, corresponding to the number of unitigs that appear in *exactly I* genomes, 1 ≤ *i* ≤ *N* . We first introduce the concept of *infix equivalent* which groups (*k* + 1)-mers based on their infix.

### Definition 1.

Infix equivalent. *Two* (*k* + 1)*-mers x, y are* infix equivalent *if x*[2 : *k*] = *y*[2 : *k*].

Therefore, a compaction between two distinct *k*-mers *x, y* happens only when the (*k* + 1)-mer *xy*[*k*] has no other infix equivalent. We call such (*k* + 1)-mers uni-mers:

### Definition 2.

Uni-mer. *A* (*k* + 1)*-mer x is a uni-mer if there is no infix equivalent y* ∈ *E*_*k*+1_(*G*), *y*≠ *x*.

Let *h*^uni-mer^(*i*) represent the frequency of uni-mers. Lastly, there is a single edge case that makes the compaction work differently: for circular unitigs, we need to cut it somewhere in its representation within the compacted de Bruijn graph in order to express it as a linear string. We call such structures *rings*, defined as:

### Definition 3.

Ring. *Given S*_*k*_(*G*), *a subset M* = {*x*_1_, …, *x*_*l*_} ⊆ *S*_*k*_(*G*) *forms a ring if consecutive k-mers overlap as x*_*i*_[2 : *k*] = *x*_*i*+1_[1 : *k* − 1] *for* 1 ≤ *i < l, and x*_*l*_[2 : *k*] = *x*_1_[1 : *k* − 1], *and no k-mer y* ∈ *S*_*k*_(*G*) *\ M overlaps with any x*_*i*_ *in either direction, i*.*e*., *y*[2 : *k*] = *x*_*i*_[1 : *k* − 1] *or x*_*i*_[2 : *k*] = *y*[1 : *k* − 1].

Let *δ*^ring^(*i*) represent the frequency of rings equal to the number of rings composed of *exactly i* genomes.

### Lemma 1.

*If in a (non-canonical) ccdBG all unitigs are uni-chromatic then the frequency of unitigs can be written as*

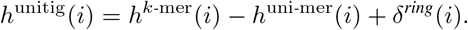

**Proof**. We first prove that the equation holds for the total number of unitigs. First, it is clear that the number of unitigs is tightly upper bounded by the number of *k*-mers and that, if we do not have rings, joining two distinct unitigs decreases the number of unitigs by one. Second, by definition two *k*-mers *x, y ∈ S*_*k*_(*G*) are joined together into a unitig if *x*[2 : *k*] = *y*[1 : *k* − 1] and there is no other *z ∈ S*_*k*_(*G*) such that either *z*[2 : *k*] = *y*[1 : *k* − 1], with *z* ≠ *x*, or *x*[2 : *k*] = *z*[1 : *k* − 1], with *z* ≠ *y*. Therefore, *xy*[*k*] is a uni-mer, because if there was a (*k* + 1)-mer *uv*[*k*] ∈ ℰ_*k*+1_(*G*) infix equivalent to *xy*[*k*] then either *u* = *z* or *v* = *z*. Since it is possible to join the ends of the same unitig, forming a ring, but we still count this as a node in ccdBG, we need to add one each time this happens. The fact that *h*^unitig^(*i*) = *h*^*k*-mer^(*i*) − *h*^uni-mer^(*i*) + *δ*^ring^(*i*) follows from the unitigs being uni-chromatic. Therefore, the color set and its frequency are the same for each *k*-mer and uni-mer in the unitig.

## 4 Interpolating unitigs

To interpolate Hill numbers for *q* ∈ {0, 1, 2} we need the expected frequency of unitigs for a given *m*, 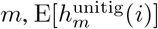. From Lemma 1 and the linearity of expectation we derive the following corollary:

### Corollary 1.

*The expected unitig frequency for* 1 ≤ *m* ≤ *N can be expressed as*

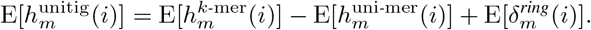

To estimate 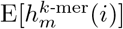, we use the hypergeometric interpolation of the observed *k*-mer frequency histogram, under the same incidence model used in (Parmigiani et al. 2024). We denote this estimator as 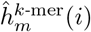:

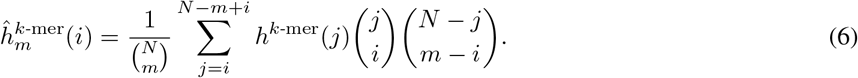

To estimate 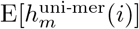 the same formula does not apply. A uni-mer is formed by taking a (*k* + 1)-mer *x* and no other infix equivalent *y* ≠ *x*. This means that (*k* + 1)-mers are not independent from each other, but they depend on the presence of other infix equivalent (*k* + 1)-mers. In order to derive a formula two observations can be made. Firstly, if a (*k* + 1)-mer is a uni-mer in the ccdBG(*G*) then it is not affected by any (*k* + 1)-mer. Secondly, if a (*k* + 1)-mer *x* is not a uni-mer, it will create a uni-mer in a subset of the genomes ℋ ⊆ *G* ccdBG(ℋ) if and only if there is no other infix equivalent *y* ≠ *x* in ℰ_*k*+1_(ℋ). This also means that a genome *G* containing two infix equivalent (*k* + 1)-mers will never form a uni-mer each time *G* ∈ ℋ.

Let *σ*(*x*) be the number of genomes *G* ∈ *G* that contain a (*k* + 1)-mer *z* ∈ ℰ_*k*+1_(*G*) with the same infix *x*[2 : *k*] = *z*[2 : *k*]. That is, *σ*(*x*) = |{*G* | *G* ∈ *G* ∧ *z* ∈ ℰ_*k*+1_(*G*) ∧ *z*[2 : *k*] = *x*[2 : *k*]|. Now suppose that a (*k* + 1)-mer *x* forms a uni-mer in exactly *c* genomes when considered individually. This means that if we take any subset of those *c* genomes, *x* will still be a uni-mer.

Let *d* = *σ*(*x*) − *c* be the number of genomes in which some (*k* + 1)-mer *y* ≠ *x* infix equivalent of *x* is present. In any subset that includes at least one genome from the *c* genomes and at least one from the *d* genomes, *x* will no longer be a unimer. This means *x* remains a uni-mer if we take any combination of genomes that includes at least one of the *c* genomes and excludes all *d* genomes. These combinations may include any of the remaining *N* − *σ*(*x*) genomes. Let *h*^infix^(*j, σ*) be the number of uni-mers that are formed in exactly *j* genomes, out of a total of *σ* genomes that contain an infix-equivalent (*k* + 1)-mer (including *x*). We count this for 1 ≤ *j* ≤ *N* and *j* ≤ *σ* ≤ *N* . Note that *h*^infix^(*σ, σ*) = *h*^uni-mer^(*σ*) since it corresponds to the number of (*k* + 1)-mers with no infix equivalent. We give an estimate for 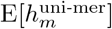 as:

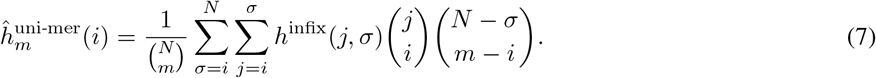

The formula counts the probability of taking a (*k* + 1)-mer *x* present in *j* genomes *i* times, and not taking any other of the *σ* genomes *m* − *i* times. Our final estimator for 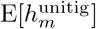 is

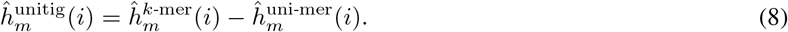

This estimator does not take into account the (*k* + 1)-mers formed by two *k*-mers exclusively present in two distinct genomes, which we refer to as *hybrid* (*k* + 1)*-mers*. Moreover, we decided to not consider E[*δ*_*m*_(*i*)] since its contribution is mostly negligible (and it requires further calculations). Finally, we can plug the estimator into the Hill number formulas, with 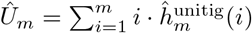. The corresponding formulas are:

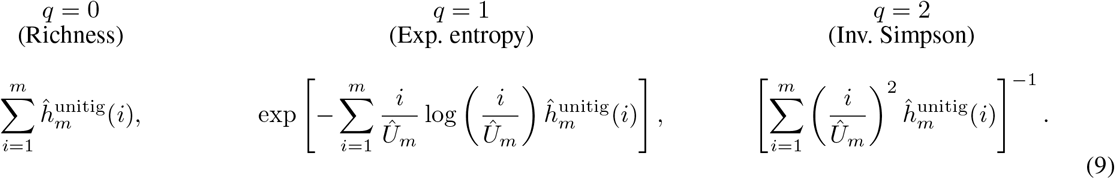

### 4.1 Computing observed frequencies from sequences

To compute the interpolation ^*q*^Δ(*m*) for 1 ≤ *m* ≤ *N*, we first require the observed pangenome frequency distributions *h*^*k*-mer^, *h*^uni-mer^, and *h*^infix^. The *k*-mer frequency *h*^*k*-mer^ can be obtained with a standard *k*-mer counter followed by counting how many *k*-mers occur *i* times as in (Parmigiani et al. 2024).

The infix and uni-mer frequencies are computed by scanning the genomes and grouping (*k* + 1)-mers by their infix using a hash table. This structure records the flanking characters of each (*k* + 1)-mer and tracks in how many genomes it forms a uni-mer while avoiding duplicate counts within the same genome. Two telomeres *a*$ and $*b* from the same genome can combine to form a (*k* + 1)-mer *ab*[*k*] when their infix match, i.e., *a*[2 : *k*] = *b*[1 : *k* − 1], even though the (*k* + 1)-mer might not appear in the genome. Conversely, a telomere *a*$ may also interact with a regular (*k* + 1)-mer *x* if *a*[2 : *k*] ≠ *x*[2 : *k*] and *a*[1] ≠ *x*[1] then no uni-mer can be formed with the infix *x*[2 : *k*] in the genome. From this structure we derive the frequency distribution *h*^infix^, which is then used to compute the expected uni-mer and unitig frequencies needed for the interpolation. A full description of the data structures, algorithms, and handling of telomeres is given in Appendix A.

Computing ^*q*^Δ(*m*) for all 1 ≤ *m* ≤ *N* requires *O*(*N* ^4^) time in the worst case, although in practice the curve is usually evaluated only for a subset of points (e.g., *s* = 30 (Hsieh et al. 2016)), reducing the complexity to *O*(*sN* ^3^).

#### Canonicity

So far the discussion considered the forward ccdBG. The same approach extends to the canonical ccdBG by orienting each (*k* + 1)-mer according to its infix. A (*k* + 1)-mer is processed in the direction where its infix is lexicographically smaller than its reverse complement, i.e., we use *x* if *x*[2 : *k*] ≤ _lex_ *x*[2 : *k*]^*rc*^ and *x*^*rc*^ otherwise. The algorithms remain unchanged except for the extraction of (*k* + 1)-mers, which must account for this orientation and for canonical telomeres. In addition, (*k* + 1)-mers whose infix is identical to its reverse complement are ignored since they cannot form uni-mers. Full implementation details are given in Appendix A.

### 4.2 Computing observed frequencies from a colored graph

Alternatively, the observed *k*-mer and infix-equivalent frequencies can be computed from a single ccdBG, assuming that we can access the color set of each unitig. A unitig of length *ℓ* with color set *C* contains *ℓ* − *k* + 1 *k*-mers and therefore contributes *ℓ* − *k* + 1 to *h*^*k*-mer^(|*C*|). Its *ℓ* − *k* internal transitions are uni-mers with the same color set and therefore contribute to *h*^infix^(|*C*|, |*C*|). These contributions can be accumulated while streaming the unitigs.

It remains to consider edges between unitigs. We store only the *k*-mers at the two extremities of each unitig and group them based on their canonical infix *x*. An edge *axb* is assigned the color set

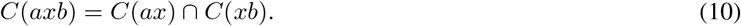

For one infix *x*, let *A*_*x*_ be the set of its 16 possible edges. Let *S*_*x*_ be the set of genomes occurring in the color set of at least one of these edges. Let *M*_*x*_ be the set of genomes occurring in the color sets of at least two distinct edges in *A*_*x*_.

Thus, *σ* = |*S*_*x*_| is the number of genomes containing at least one infix equivalent. A genome in *M*_*x*_ contains more than one distinct infix equivalent and therefore cannot contribute to any possible uni-mer with infix *x*. Consequently, for each candidate edge *e* ∈ *A*_*x*_, only the genomes containing that edge and no other candidate are counted *j*_*e*_ = |*C*(*e*) *\ M*_*x*_|.

For every candidate with *j*_*e*_ *>* 0, we increment *h*^infix^(*j*_*e*_, *σ*). Grouping extremities by infix avoids storing all (*k* + 1)-mers. No separate telomere correction is needed because sequence ends are already reflected in the graph topology.

### 4.3 Approximating interpolated frequencies

The exact formula for 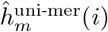 requires *O*(*N* ^3^) time for each sample size *m*. Although this cost does not depend on genome length, unlike repeatedly constructing the graph by sampling, it becomes expensive when *N* is large. We therefore approximate both the *k*-mer and uni-mer frequencies using a Bernoulli sampling model.

Consider an entry *h*^infix^(*j, σ*). The candidate (*k* + 1)-mer forms a uni-mer in *j* genomes, while *σ* is the total number of genomes containing a (*k* + 1)-mer with the same infix. Thus, the remaining *σ* − *j* genomes contain an infix-equivalent competitor.

For a fixed sample size *m*, the candidate contributes to 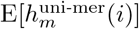 if exactly *i* of the *j* genomes containing the candidate are selected, and none of the genomes containing an infix-equivalent competitor are selected. The remaining *m* − *i* genomes must therefore be chosen from the *N* − *σ* genomes that contain neither the candidate nor one of its competitors. This gives the exact contribution

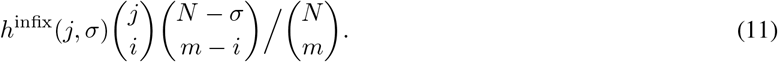

To reduce the computational complexity, we use a Bernoulli model where each of the *N* genomes is included with probability *p* = *m/N*, independently of the others. Thus, the number of included genomes is not fixed to *m*, but has expected value *m*. We do not use this model to actually sample random subsets; we use it only to approximate the expected value of each histogram entry 1 ≤ *i* ≤ *m*.

Under this approximation, the candidate contributes to frequency *i* when exactly *i* of the *j* genomes containing it are included, while all other *σ* − *i* genomes containing either the same candidate or an infix-equivalent competitor are excluded, therefore

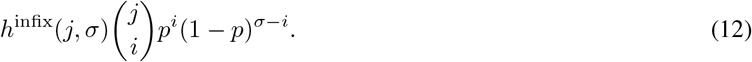

The approximate uni-mer frequency is

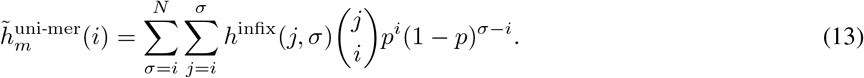

Since (1 − *p*)^*σ*−*i*^ = (1 − *p*)^*j*−*i*^(1 − *p*)^*σ*−*j*^, and since the indices satisfy *i* ≤ *j* ≤ *σ* ≤ *N*, we can invert the indices *j* and *σ*:

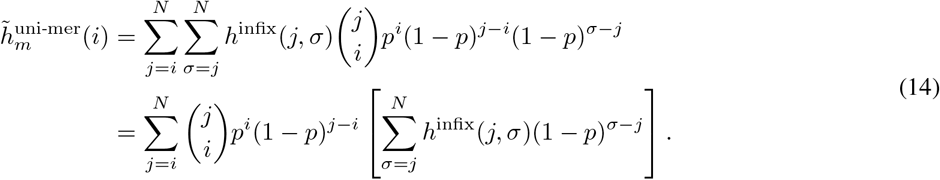

For a fixed *m*, all values *j* in the inner sum of Equation 14 require *O*(*N* ^2^) time, and all frequencies 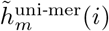 require another *O*(*N* ^2^) time. The total complexity for *s* sample sizes is therefore reduced to *O*(*sN* ^2^).

We apply the same Bernoulli sampling model to the *k*-mer frequency,

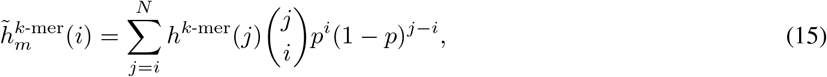

and compute the approximate unitig frequency as

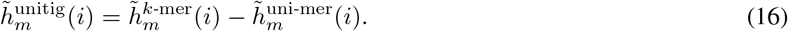

The Bernoulli approximation is especially useful for reducing the cost of the full histogram, but it can be less accurate for values of *i* close to *m*. These entries correspond to items that are present in almost all selected genomes. Since the Bernoulli model does not condition on selecting exactly *m* genomes, it can distort these entries.

For this reason, we use a hybrid estimator. Given a value *r*, the last *r* frequencies, max(1, *m* − *r* + 1) ≤ *i* ≤ *m*, are computed with the exact formulas for both *k*-mers and uni-mers, while the remaining frequencies use the Bernoulli approximation. Setting *r* = 0 gives the pure Bernoulli approximation, whereas *r* ≥ *m* gives the hypergeometric estimator.

The exact tail can be computed jointly for all sample sizes. Let *d* = *m* − *i* be the offset from the end of the histogram and define

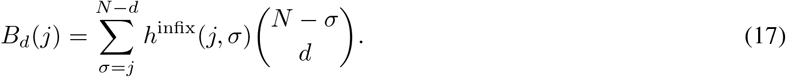

Then

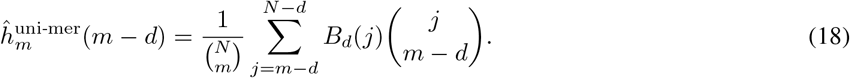

For each of the *r* offsets, all values *B*_*d*_(*j*) require *O*(*N* ^2^) time, after which each sampled *m* requires *O*(*N*) time. The exact tail therefore costs *O*(*rN* ^2^ + *srN*). Together with the Bernoulli approximation, the total complexity is *O*((*s* + *r*)*N* ^2^).

## 5 Extrapolating unitigs

To extrapolate the Hill numbers ^*q*^Δ(*N* + *m*^∗^), for *m*^∗^ ≥ 1, we need an estimator 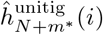 that can be used in any of the formulas in Equation (9). Similarly to Equation (8) for the interpolation, we propose an estimator 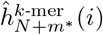 and an estimator 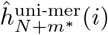. To construct them, we consider the three processes that contribute to the growth of a pangenome graph. The first two are the addition of new, previously unseen *k*-mers and uni-mers. The third is the process of uni-mers being broken into *k*-mers by the addition of *k*-mers which form new infix equivalents in ℰ_*k*+1_ (see Appendix C for the theoretical formula).

First, we define an estimator 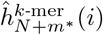. Suppose we want to estimate the histogram at step *N* + 1. The value in *h*_*N*+1_(*i*) can increase only if some *k*-mers present in *h*(*i* − 1) were found in the (*N* + 1)th genome. Let this occur with probability *p*_*i*−1_. Likewise, *h*_*N*+1_(*i*) can decrease only if some *k*-mers in *h*(*i*) are found in the (*N* + 1)th genome. Since the true value of *p*_*i*_ is unknown, we approximate it with 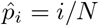. Therefore, for each *i >* 1, the expected number of *k*-mers is given by:

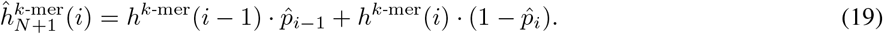

For *i* = 1, we need to account for both observed singleton *k*-mers that remain singletons and previously unseen *k*-mers introduced by the (*N* + 1)th genome. We estimate *ĥ*^*k*-mer^(0), the number of *k*-mers in the pangenome that have not yet been observed, using the Chao2 estimator (Gotelli and Chao 2013), and define the probability that an individual previously unseen *k*-mer is found when adding one new genome as 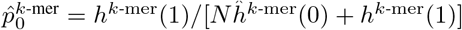. Therefore,

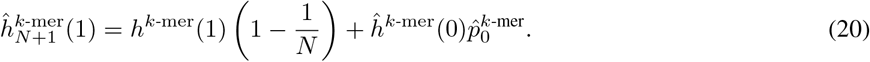

We can extend this to *m*^∗^ additional genomes using the binomial distribution for *i >* 1:

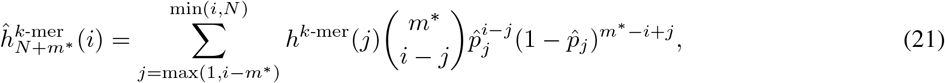

which gives the expected number of *k*-mers that were previously seen *j* times and are now seen *i* times, meaning they appeared exactly *i* − *j* times in the new *m*^∗^ genomes. This also includes the *k*-mers that were already observed *i* times and did not appear in any of the *m*^∗^ new genomes (the *j* = *i* case). For *i* = 1, we use

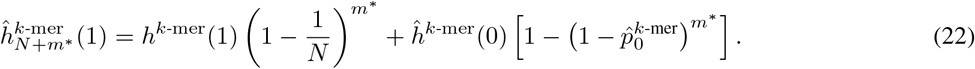

We now focus on defining an estimator for the uni-mer histogram 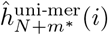. We follow a similar approach as 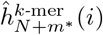, where the value of 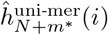 changes based on the value of the histogram in positions *j* ≤ *i*. Uni-mers that were present in exactly *i* − *m*^∗^ genomes can appear in *i* genomes if and only if they are found in all *m*^∗^ of the new genomes. Similarly, uni-mers previously present in *i* − *m*^∗^ + 1 genomes may appear in *i* genomes if and only if they are found in *m*^∗^ − 1 of the new genomes, and so on. Finally, uni-mers that were already present in *i* genomes remain at count *i* if they are not found in any of the *m*^∗^ new genomes.

On top of this, we must account for the possibility that a uni-mer breaks due to the appearance of a new infix equivalent. Let this occur with probability *ρ*. To approximate *ρ* we define:

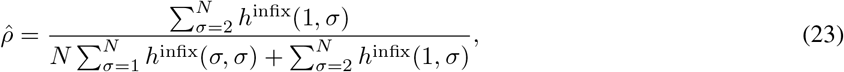

where the diagonal entries count observed uni-mers, since *h*^infix^(*σ, σ*) = *h*^uni-mer^(*σ*), and *h*^infix^(1, *σ*) counts singleton competing (*k* + 1)-mers within infix-equivalence classes spanning *σ* ≥ 2 genomes. For *i >* 1, the estimator is given by:

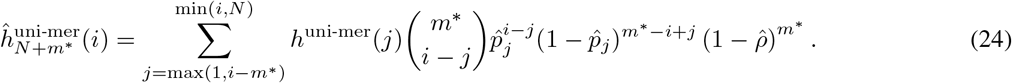

For *i* = 1, let 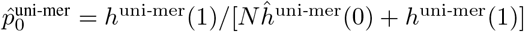. We use

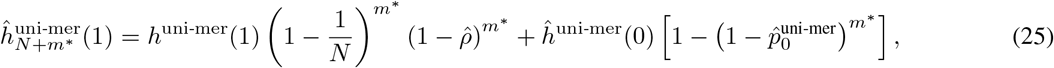

where *ĥ*^uni-mer^(0) is the estimated number of uni-mers in the pangenome that have not yet been seen. This is computed analogously to *ĥ*^*k*-mer^(0), but substituting *h*^*k*-mer^(1) and *h*^*k*-mer^(2) with *h*^uni-mer^(1) and *h*^uni-mer^(2), respectively. For *m*^∗^ *>* 1, these singleton-bin estimators approximate every previously unseen item discovered at least once as occurring in exactly one of the additional genomes.

## 6 Benchmarks

We evaluated Pangrowth through six experiments. First, we validated the hypergeometric interpolation estimator against an exhaustive-subset reference. Second, we compared its runtime and memory usage with repeated construction of colored compacted de Bruijn graphs using Bifrost and GGCAT. Third, we evaluated the graph-based mode, where Pangrowth receives an already constructed ccdBG and extracts the observed frequencies from its color information. Fourth, we tested the Bernoulli-hybrid approximation against the exact hypergeometric interpolation. Fifth, we evaluated extrapolation by predicting the complete dataset from held-out genomes. Finally, we tested the sensitivity of the Hill-number curves to the graph order *k*.

The runtime and memory comparison is not a one-to-one comparison of identical tasks: constructing a ccdBG and estimating its interpolated or extrapolated size are different tasks. Nevertheless, repeated graph construction is the only direct empirical baseline available, and it provides a useful reference for the computational cost of our method.

The two approaches are not mutually exclusive. Our method can also be applied to an existing ccdBG by taking the graph as input and extracting the color information, as described in Subsection 4.2. We use this mode for the human pangenome experiment below.

We selected *k* using a heuristic similar to that of Sheikhizadeh Anari et al. (2018). Given the genome length, we first found the smallest *k* for which the probability that a random *k*-mer occurs in the genome is below 10^−4^. This gave 18 for *E. coli* and 21 for *A. thaliana*. We rounded upward to odd values to exclude reverse-complement palindromic *k*-mers, then used one additional odd step as a conservative margin, giving *k* = 21 and *k* = 23, respectively. We also computed E[*h*^unitig^] on a subset of 8 *E. coli* genomes by constructing the graph for every possible genome subset ℋ ⊆ *G* and averaging the results. All computations ran on a server with 28-core Intel Xeon CPUs (2.6 GHz) and 64 GB RAM.

We refer to an estimator by both its observed-frequency source and its interpolation: sequence-based or graph-based, followed by hypergeometric or Bernoulli-hybrid. Unless stated otherwise, the experiments below use the sequence-based hypergeometric estimator with telomeres.

### 6.1 Interpolation

#### Hypergeometric estimator

Firstly, we validate the sequence-based hypergeometric estimator for each *m*, 1 ≤ *m* ≤ 8, using 8 *E. coli* genomes (Appendix B, Table 2). We evaluate two variants: one counting only (*k* + 1)-mers and one that also includes telomeres. Ignoring telomeres slightly reduces memory usage. Even without them, the estimates are very close to the exhaustive-subset reference, while including telomeres matches it here because no rings or hybrid (*k* + 1)-mers occur. In the rest of these experiments we use the version with telomeres.

Secondly, we compare its Hill numbers with the exhaustive-subset reference (Appendix B, Table 3). The values are very close; small differences arise from assuming uni-chromatic unitigs, whereas some unitigs are heterochromatic. The discrepancy is negligible and is illustrated in Figure 1.

**Figure 1.**
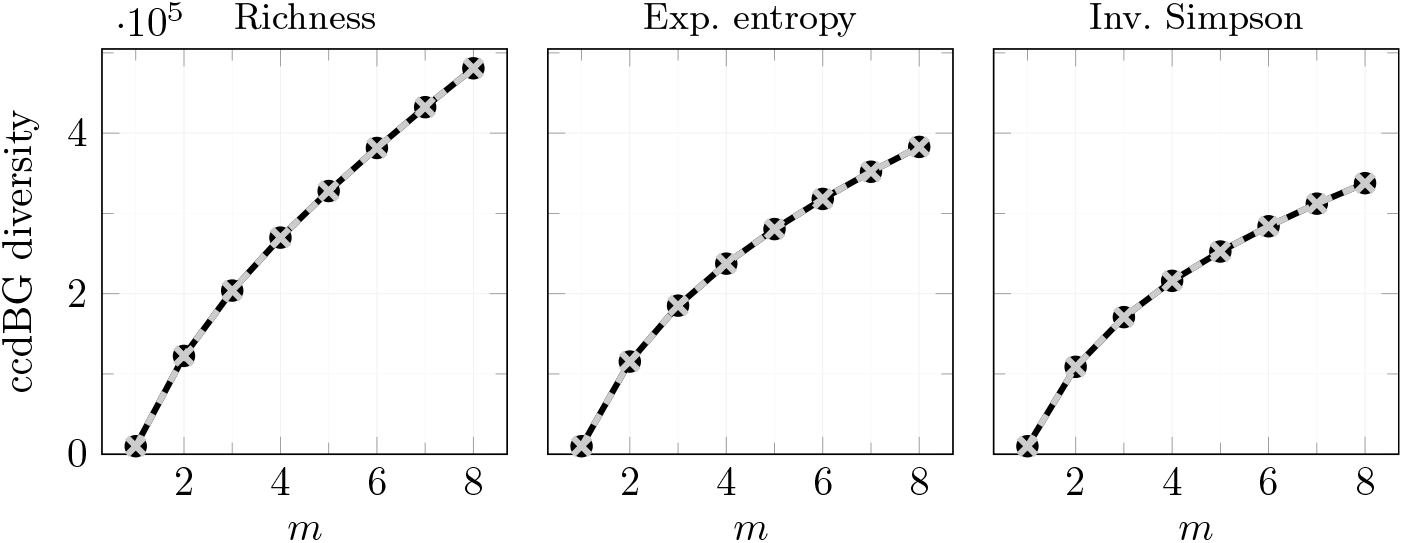
Interpolated Hill numbers for different *m* values for 8 *E. coli* genomes. The solid black dots represent the exhaustive-subset reference. The gray crosses represent the sequence-based hypergeometric estimates.

#### Comparison with sampling graph construction

Next, we place the time and memory usage of the sequence-based hypergeometric estimator in the context of repeated graph construction. Several tools can build compacted de Bruijn graphs, including Bcalm2 (Chikhi et al. 2016), Bifrost (Holley and Melsted 2020), Cuttlefish (Khan et al. 2022), Cdbgtricks (Hannoush et al. 2024), GGCAT (Cracco and Tomescu 2023), Nexus (Depuydt et al. 2023), and TwoPaCo (Minkin et al. 2016). Among these, Bifrost, GGCAT, and Nexus support colored compacted de Bruijn graphs. We used Bifrost and GGCAT for the repeated-construction benchmark.

Bifrost supports updating the graph with batches of new genomes, and we used it in this mode. In contrast, GGCAT does not support incremental updates, so it was re-run from scratch for each subsample. We benchmarked Bifrost (v2.8.12), GGCAT (v2.0.0), and our tool Pangrowth on two datasets: one composed of 1000 *Escherichia coli* genomes and another composed of 20 *Arabidopsis thaliana* genomes. Our tool was run once on each whole dataset. Details are given in Appendix B.

Table 1 reports the corresponding time and space usage. On the *E. coli* dataset, the sequence-based estimator took 15.19 minutes. One repeated-construction series took 24.25 minutes with Bifrost and 5.28 minutes with GGCAT; ten series took 242.53 and 52.85 minutes, respectively. On the *A. thaliana* dataset, the corresponding times were 1.61 minutes for Pangrowth, 52.66 minutes for one Bifrost series, and 4.65 minutes for one GGCAT series.

**Table 1.** Time (wall clock) and space usage (Maximum Resident Set Size) for the sequence-based hypergeometric estimator and repeated graph construction on 1000 *E. coli* genomes and 20 *A. thaliana* genomes.

| program | *E. coli* ($N = 1000$) | | *A. thaliana* ($N = 20$) | |
| --- | --- | --- | --- | --- |
|  | time [m] | space [GB] | time [m] | space [GB] |
| Pangrowth, sequence-based (total) | 15.19 | 9.30 | 1.61 | 21.31 |
| Histogram $k$-mer | 3.25 | 1.81 | 0.78 | 8.74 |
| Histogram infix eq. | 2.97 | 9.30 | 0.83 | 21.31 |
| Hill numbers | 8.97 | 0.01 | 0.0003 | 0.004 |
| Bifrost update (10 sampling) | 242.53 | 3.17 | 526.63 | 7.48 |
| Bifrost update (1 sampling) | 24.25 | 3.17 | 52.66 | 7.48 |
| GGCAT build (10 sampling) | 52.85 | 2.15 | 46.48 | 2.53 |
| GGCAT build (1 sampling) | 5.28 | 2.15 | 4.65 | 2.53 |

**Table 2.** Interpolated richness values for different *m* computed with the sequence-based hypergeometric estimator, with (w/) and without (w/o) telomeres, versus the exhaustive-subset reference for 8 *E. coli* genomes.

| $m$ | w/o telomeres | w/ telomeres | exhaustive |
| --- | --- | --- | --- |
| 1 | 9997.38 | 10 003.50 | 10 003.50 |
| 2 | 122 496.36 | 122 503.20 | 122 503.20 |
| 3 | 203 876.64 | 203 884.30 | 203 884.30 |
| 4 | 269 846.37 | 269 854.70 | 269 854.70 |
| 5 | 327 885.70 | 327 894.70 | 327 894.70 |
| 6 | 381 515.04 | 381 524.80 | 381 524.80 |
| 7 | 432 353.50 | 432 363.90 | 432 363.90 |
| 8 | 480 795.00 | 480 806.00 | 480 806.00 |

**Table 3.**
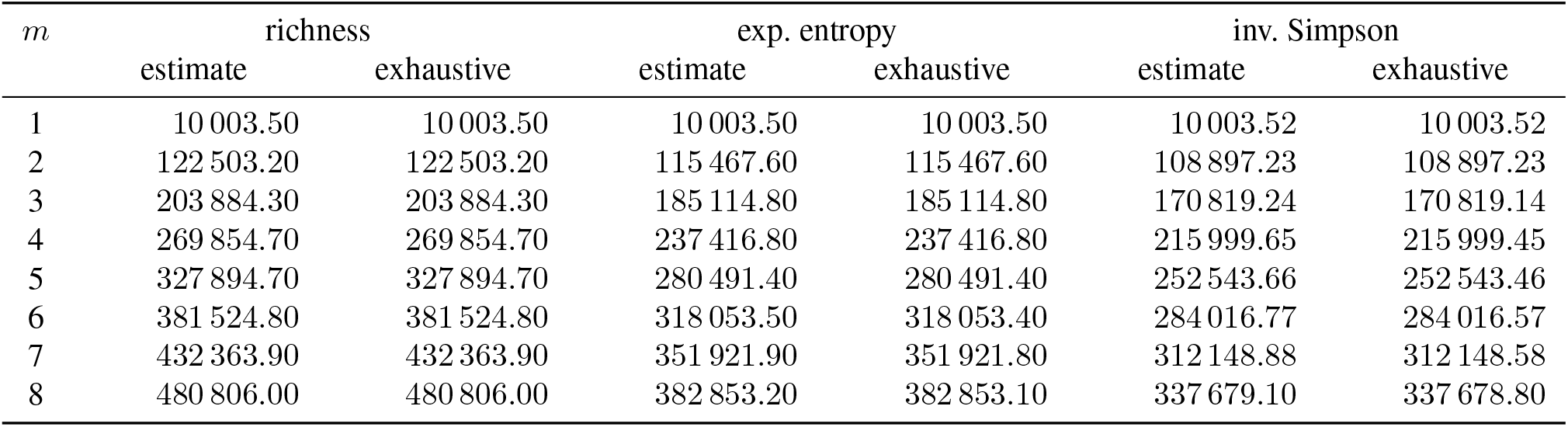
Comparison of Hill numbers interpolated with the sequence-based hypergeometric estimator versus the exhaustive-subset reference for 8 *E. coli* genomes.

Peak memory is also reported only to indicate the resources required by each tool, since graph construction and histogram-based estimation produce different outputs and make different use of disk storage. For Pangrowth, the largest contribution was the construction of the infix-equivalent histogram.

Lastly, we compared the sequence-based hypergeometric estimates with Hill numbers obtained from repeated Bifrost graph construction. Figures 8 and 9 in Appendix B show the two datasets respectively. For Bifrost, the unitig histograms were averaged over 10 random genome subsets for each *m*. On the *E. coli* dataset, the estimates closely match. On the *A. thaliana* dataset, they show the same trend but fluctuate more, suggesting that 10 repetitions were not enough to obtain a stable average. Since the Hill numbers from the graphs constructed from Bifrost and GGCAT are the same we only show the one from Bifrost.

#### Graph-based frequencies from Human pangenome

The current sequence-based implementation has high RAM requirements on large genomes. We therefore used the graph-based procedure instead: we first constructed the ccdBG with GGCAT and then extracted the observed frequencies directly from the graph colors. We constructed the graph with GGCAT using *k* = 31, a common value for human genomes (Lee et al. 2020). We then used the C++ API of GGCAT to read the graph in Pangrowth and compute the observed frequencies from the color information.

Overall, the process took 90.64 minutes. Most of the time was spent constructing the graph, which took 87.38 minutes, while the Pangrowth step took 3.26 minutes. This shows that, once the graph is available, extracting the frequencies and computing the Hill numbers adds only a small overhead.

The maximum resident memory was 12.34 GB for the GGCAT step and 6.89 GB for the Pangrowth step. However, GGCAT also required 280.66 GB of disk space during graph construction.

Figure 10 and Table 4 in Appendix B.1 show the Hill numbers over *m* for 100 human genomes and summarize the benchmark results, respectively.

**Table 4.** Time and maximum resident memory for a 100 human genomes.

|  | time [m] | space [GB] |
| --- | --- | --- |
| GGCAT build | 87.38 | 12.34 |
| Pangrowth, graph-based (total) | 3.26 | 6.89 |
| Histogram $k$-mer+infix eq. | 3.25 | 6.89 |
| Hill numbers | 0.01 | 0.01 |

#### Bernoulli-hybrid approximation

We evaluate the approximation introduced in Section 4.3 on the 1000 genome *E. coli*, 20 genome *A. thaliana*, and 100 genome human datasets. Figure 2 shows that the Bernoulli-hybrid and hypergeometric estimates closely overlap for all three Hill numbers.

**Figure 2.**
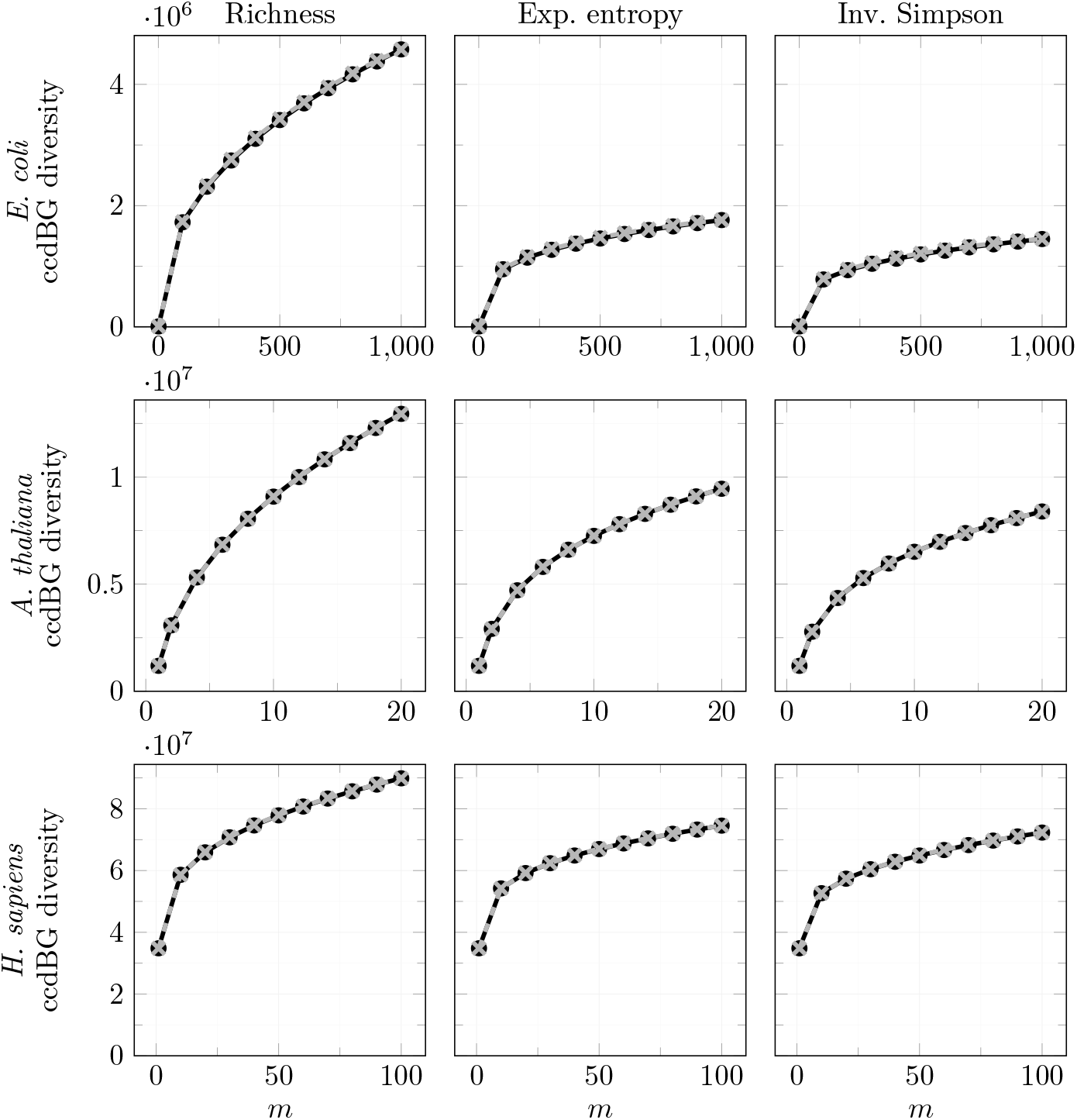
Hill numbers interpolated with the hypergeometric estimator (black circles) and the Bernoulli-hybrid estimator (gray dashed lines and crosses) for 1000 *E. coli* genomes, 20 *A. thaliana* genomes, and 100 human genomes (top to bottom). The maximum possible number of corrections was set to *R* = 20 and tolerance *τ* = 0.001. Columns show richness, exponential entropy, and inverse Simpson index.

We use an adaptive rule to decide how many high-frequency histogram entries are corrected. We set a maximum number of corrections *R* = 20 and a tolerance *τ* = 0.001. For each sample size *m*, we start from the last histogram entry *i* = *m* and move downwards. At each step, we replace the Bernoulli value with the exact hypergeometric value, update the three Hill numbers, and compare them with the values obtained before the replacement.

A correction is considered stable if the relative change is at most *τ* for all three Hill numbers: richness, exponential entropy, and inverse Simpson index. We stop after three consecutive stable corrections, or when the maximum tail length *R* is reached.

The stopping condition controls the effect of changing each additional histogram value, but does not guarantee a bounded difference from the hypergeometric result. In particular, the errors of the *k*-mer and uni-mer estimates can cancel, so the final error does not need to decrease monotonically with the number of changed values.

For *E. coli*, the procedure changed at most 14 histogram values across the ten interpolation points, with a mean of 11.20. For *A. thaliana*, it changed at most 11 values (mean 7.30), and for the human pangenome it changed at most 19 values (mean 14.80). No interpolation point reached the cap in any dataset. Thus, in these experiments, a larger value of *R* would provide no benefit, when used with this stopping rule. Empirically, the maximum relative error across the three Hill numbers and all interpolation points was 1.72% for *E. coli*, 0.068% for *A. thaliana*, and 0.164% for the human pangenome (Appendix B.2, Figure 11).

The adaptive interpolation took 0.135 seconds for *E. coli*, 0.0018 seconds for *A. thaliana*, and 0.0030 seconds for the human pangenome. The exact interpolation instead required 464.39, 0.0087, and 0.54 seconds, respectively. The adaptive method was therefore approximately 3435, 4.9, and 178 times faster. Results for fixed values *r* ∈ {0, 1, 3, 5, 10} are also reported in Table 5 in Appendix B.2.

**Table 5.** Maximum relative error over the sampled values of *m* and runtime of fixed-tail and adaptive Bernoulli-hybrid interpolation. Errors are percentages relative to hypergeometric interpolation. The adaptive method uses *R* = 20 and *τ* = 0.001.

| dataset | $r$ | richness | exp. entropy | inv. Simpson | time [s] |
| --- | --- | --- | --- | --- | --- |
| *E. coli* ($N = 1000$) | 0 | 277.190 | 277.190 | 277.190 | 0.114 |
|  | 1 | 2.248 | 2.787 | 3.568 | 0.127 |
|  | 3 | 0.637 | 1.101 | 1.404 | 0.124 |
|  | 5 | 0.358 | 0.729 | 0.898 | 0.128 |
|  | 10 | 0.728 | 1.317 | 1.686 | 0.147 |
|  | adaptive | 0.742 | 1.339 | 1.719 | 0.135 |
|  | exact | — | — | — | 464.39 |
| *A. thaliana* ($N = 20$) | 0 | 34.393 | 34.668 | 35.268 | 0.0017 |
|  | 1 | 8.217 | 7.931 | 7.545 | 0.0020 |
|  | 3 | 3.161 | 3.974 | 4.297 | 0.0019 |
|  | 5 | 1.076 | 1.306 | 1.340 | 0.0020 |
|  | 10 | 0.057 | 0.061 | 0.050 | 0.0021 |
|  | adaptive | 0.063 | 0.068 | 0.056 | 0.0018 |
|  | exact | — | — | — | 0.0087 |
| *H.sapiens* ($N = 100$) | adaptive | 0.163 | 0.146 | 0.135 | 0.0030 |
|  | exact | — | — | — | 0.5357 |

### 6.2 Extrapolation

We evaluated extrapolation from 25%, 50%, and 75% of the genomes to the complete dataset size for both *E. coli* and *A. thaliana*. For each fraction, we generated 10 random subsets by permuting the genomes and taking the corresponding prefix; the remaining genomes were held out. Figure 3 illustrates the Hill-number curves obtained from the 50% subsets.

**Figure 3.**
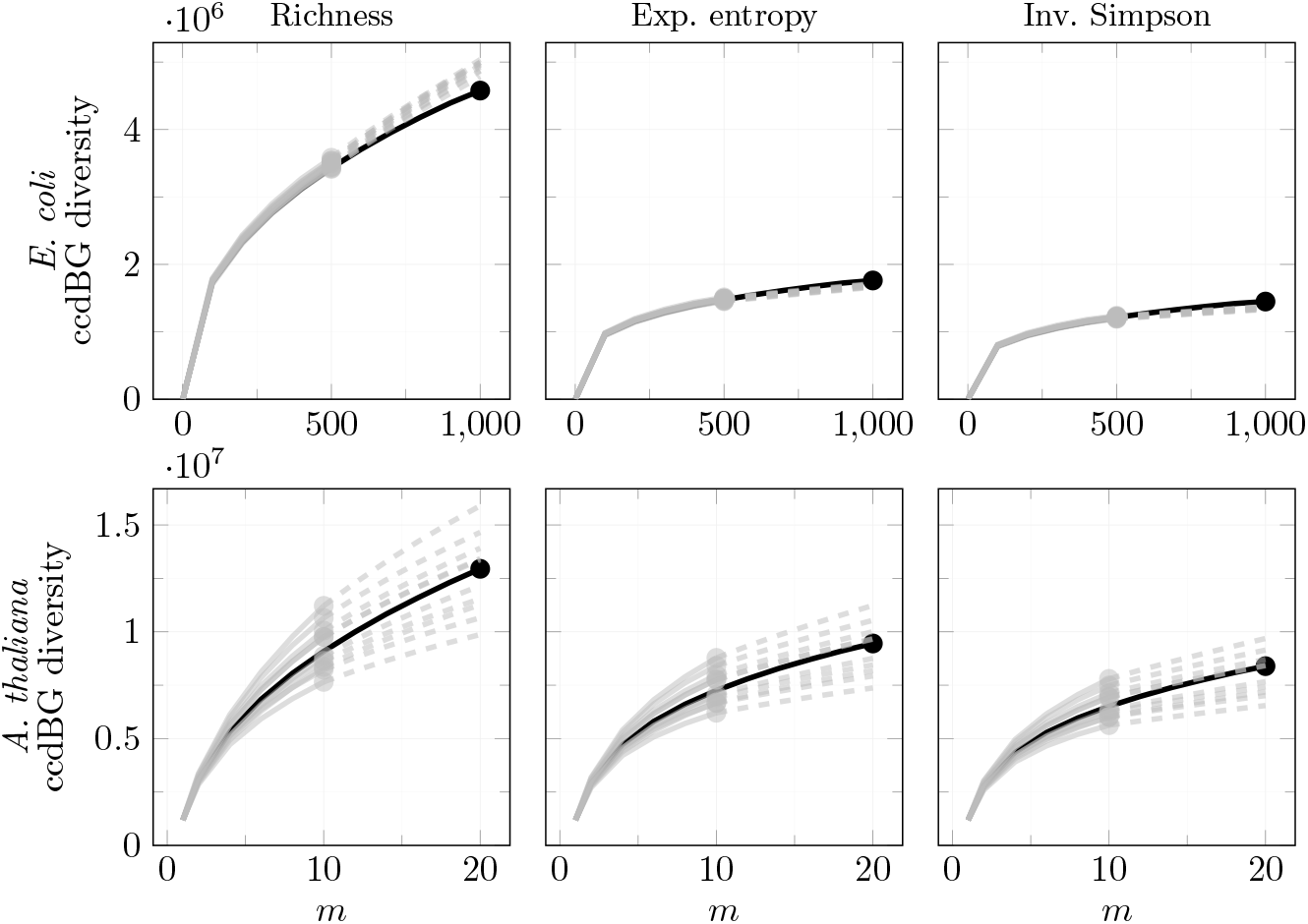
Hill numbers from the complete datasets (black) and 10 randomized 50% subsets (gray) for 1000 *E. coli* genomes (top) and 20 *A. thaliana* genomes (bottom). Solid gray lines show interpolation up to the observed subset size and dashed gray lines show extrapolation to the complete dataset size. Columns show richness, exponential entropy, and inverse Simpson index.

The extrapolations follow the full-data curves, with greater variability among the *A. thaliana* subsets. The signed relative errors at the complete dataset size for all three observed fractions are shown in Figure 12 in Appendix C.

As expected, the errors decrease as the observed fraction increases. Across the tested fractions, relative errors decrease from about 20–30% to about 5%. The remaining bias depends on how representative the observed subset is. This effect is limited for *E. coli*, where the estimates remain relatively stable across subsets, but it is more pronounced for *A. thaliana*, where the extrapolation depends more strongly on which genomes are included in the subset.

#### Sensitivity of Hill numbers to *k*

We tested the sensitivity of the Hill numbers to the graph order using *k* ∈ {19, 21, 23, 25, 27, 29} for both *E. coli* and *A. thaliana*. Figure 4 shows the three Hill-number curves for each value of *k*.

**Figure 4.**
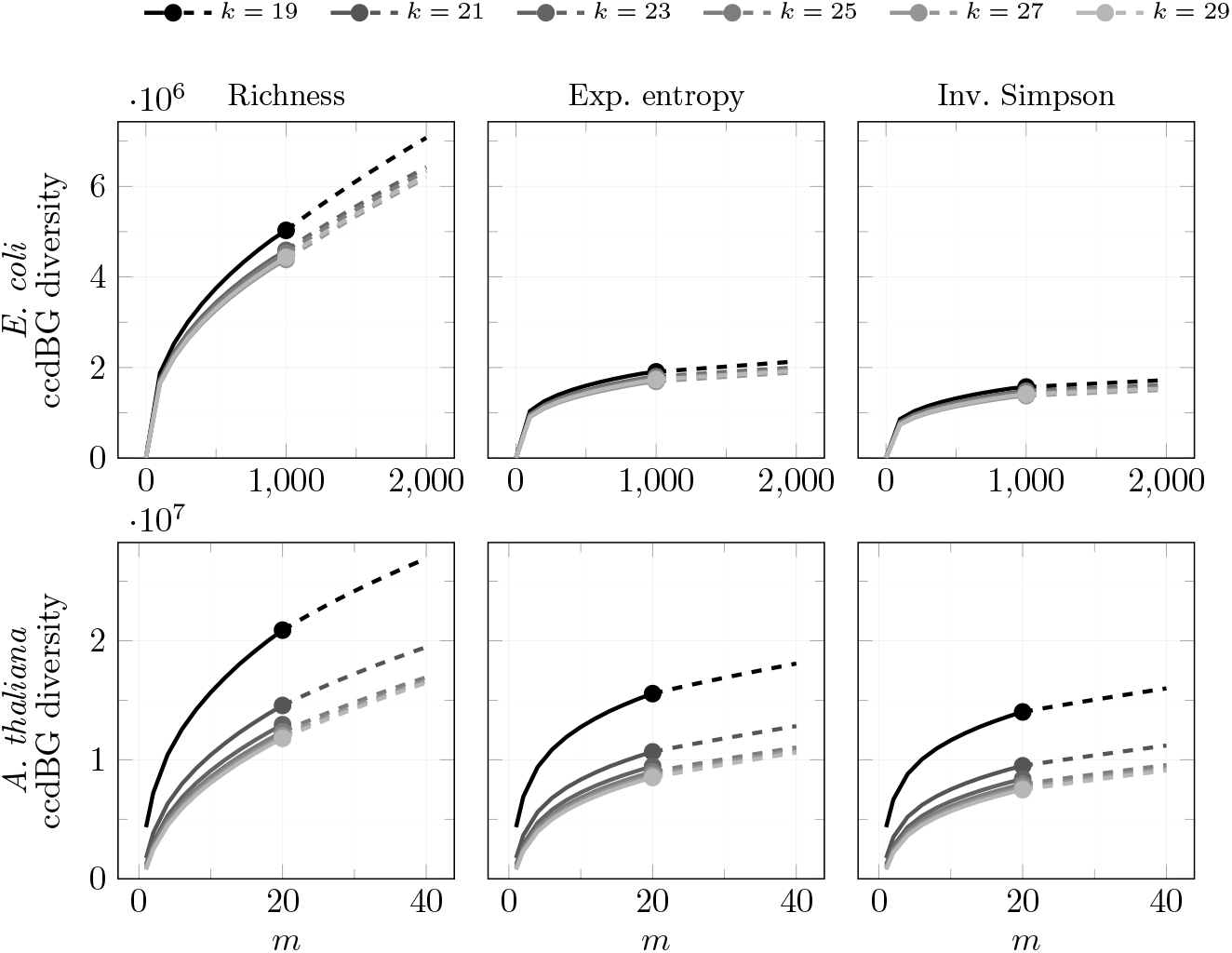
Sensitivity of the Hill-number curves to *k* for 1000 *E. coli* genomes (top) and 20 *A. thaliana* genomes (bottom). Columns show richness, exponential entropy, and inverse Simpson index. Dots mark the observed values. The values used in the main experiments were *k* = 21 and *k* = 23, respectively.

As *k* increases, the Hill numbers tend to decrease. This is expected because larger *k*-mers are less likely to overlap by chance, and therefore unitigs are less likely to be split by spurious branching. The curves stabilize around the values used in the other experiments, namely *k* = 21 for *E. coli* and *k* = 23 for *A. thaliana*.

## 7 Pangenome graph comparison

We used our method to compare the Hill numbers of the ccdBG graphs for 12 bacterial species from (Parmigiani et al. 2024). Because species have different genome lengths, this factor must be considered when comparing diversity: species with longer genomes may appear more diverse simply because more nucleotides are sampled. This issue is well known in ecology, where comparing communities from areas with different sampling efforts can give misleading results. Normalizing by genome size is not sufficient, as dividing by genome length implicitly assumes that halving the genome halves the diversity, which is generally not true.

A better approach is to use the concept of *coverage*, which quantifies how completely a sample captures the total diversity. A pangenome can contain many shared items and a long tail of unique, rare items. If we sequence a few genomes, we may observe mostly common items, while many rare ones remain undetected. Coverage is the fraction of the items (counted with multiplicity) that is represented in the sample. Low coverage means the observed diversity underestimates the true diversity, especially for richness, which is sensitive to rare elements.

We define the coverage for *k*-mers, and use it to compare two ccdBGs with the same *k*-mer coverage. Due to the *k*-mer distribution (Equation (1)) we can use the estimators defined by (Chao and Jost 2012). Let *Ĉ*_*m*_ be the coverage estimator for *m* genomes. Formulas are reported in Appendix E.

The number and length distribution of the unitigs in each complete graph are reported in Table 6 (Appendix D.1). These statistics provide the scale of the graphs underlying the diversity.

**Table 6.**
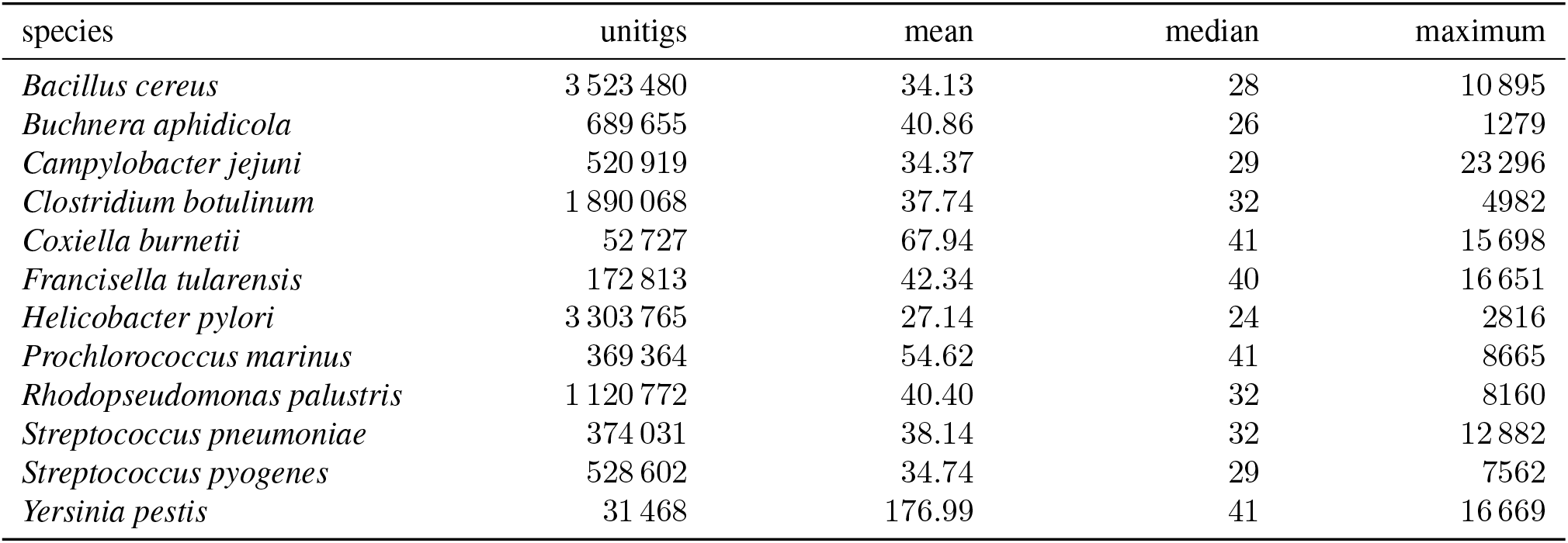
Number and length distribution (in bp) of unitigs in the complete pangenome graph (*k* = 21) for each bacterial species.

**Table 7.** Heaps-law exponents and adjusted *R*^2^ for *k*-mer and ccdBG richness for the 12 bacterial species. Species marked with ∗ have a negative ccdBG *α* and are excluded from Figure 7.

| species | $N$ | $\alpha_{k\text{-mer}}$ | $\alpha_{\text{ccdBG}}$ | $R^2_{k\text{-mer}}$ | $R^2_{\text{ccdBG}}$ |
| --- | --- | --- | --- | --- | --- |
| *Bacillus cereus* | 83 | 0.761 | 0.630 | 0.997 | 0.975 |
| *Buchnera aphidicola* * | 35 | 0.227 | -0.414 | 0.979 | 0.849 |
| *Campylobacter jejuni* | 234 | 0.856 | 0.832 | 0.998 | 0.996 |
| *Clostridium botulinum* | 50 | 0.692 | 0.399 | 0.986 | 0.974 |
| *Coxiella burnetii* | 13 | 0.909 | 0.955 | 0.978 | 0.991 |
| *Francisella tularensis* | 57 | 0.766 | 0.882 | 0.996 | 0.996 |
| *Helicobacter pylori* | 234 | 0.647 | 0.501 | 1.000 | 0.986 |
| *Prochlorococcus marinus* * | 9 | 0.215 | -0.837 | 0.966 | 0.954 |
| *Rhodopseudomonas palustris* | 8 | 0.510 | 0.172 | 0.997 | 0.608 |
| *Streptococcus pneumoniae* | 88 | 0.737 | 0.702 | 0.994 | 0.991 |
| *Streptococcus pyogenes* | 247 | 0.736 | 0.704 | 0.999 | 0.997 |
| *Yersinia pestis* | 56 | 0.838 | 0.724 | 0.916 | 0.984 |

For visualization purposes, we divided the 12 species into two groups of six each: a first group with the species with more than one billion distinct *k*-mers in their pangenome (*B. cereus, B. aphidicola, C. botulinum, H. pylori, P. marinus, R. palustris*) and a second group with species with at most one billion *k*-mers (*C. jejuni, C. burnetii, F. tularensis, S. pneumoniae, S. pyogenes, Y. pestis*). Figures 5 and 6 show the Hill numbers of ccdBG and *k*-mer for the first and second group, respectively.

**Figure 5.**
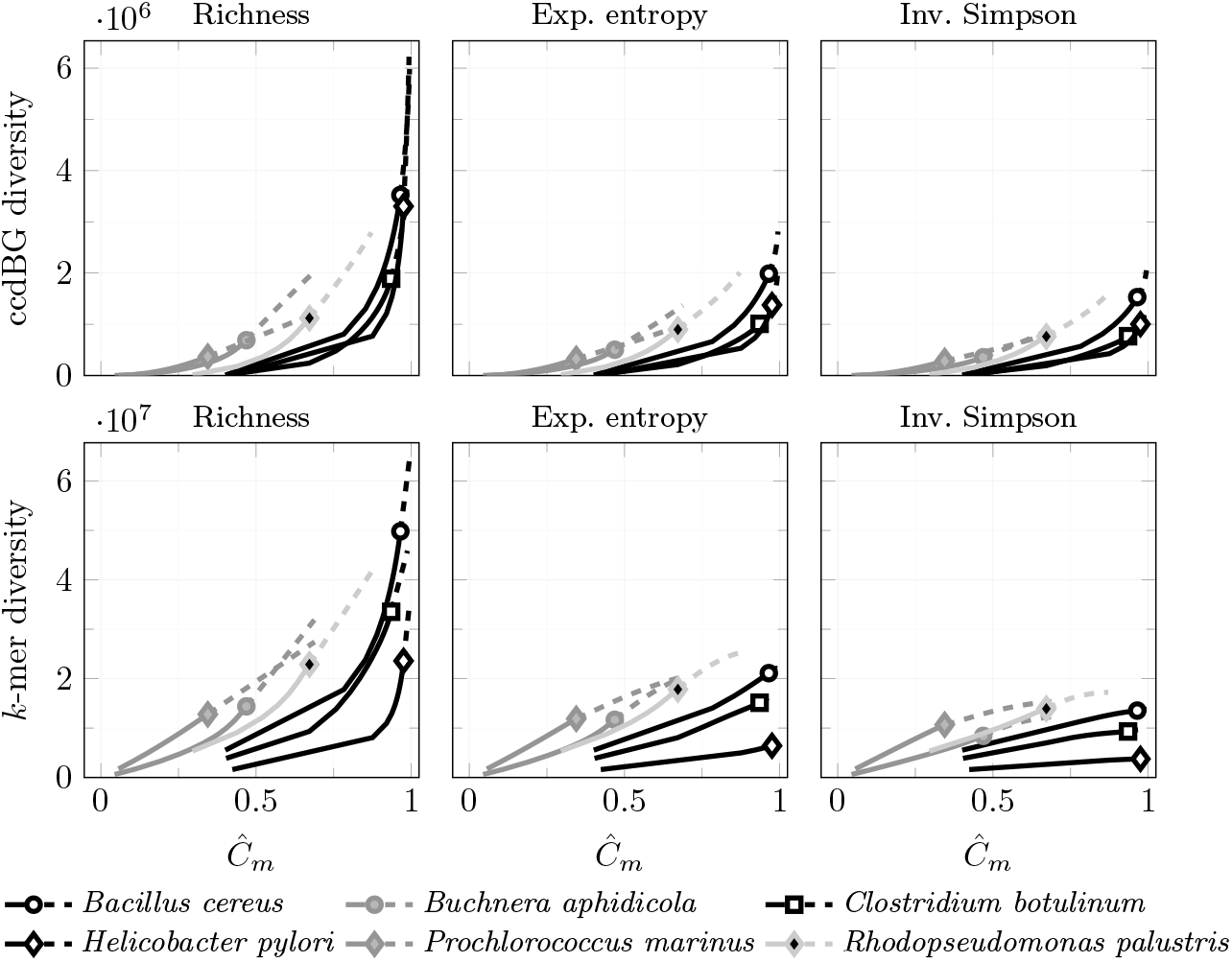
Hill numbers of ccdBG and *k*-mers for the group of species with more than 1 billion distinct *k*-mers over coverage.

**Figure 6.**
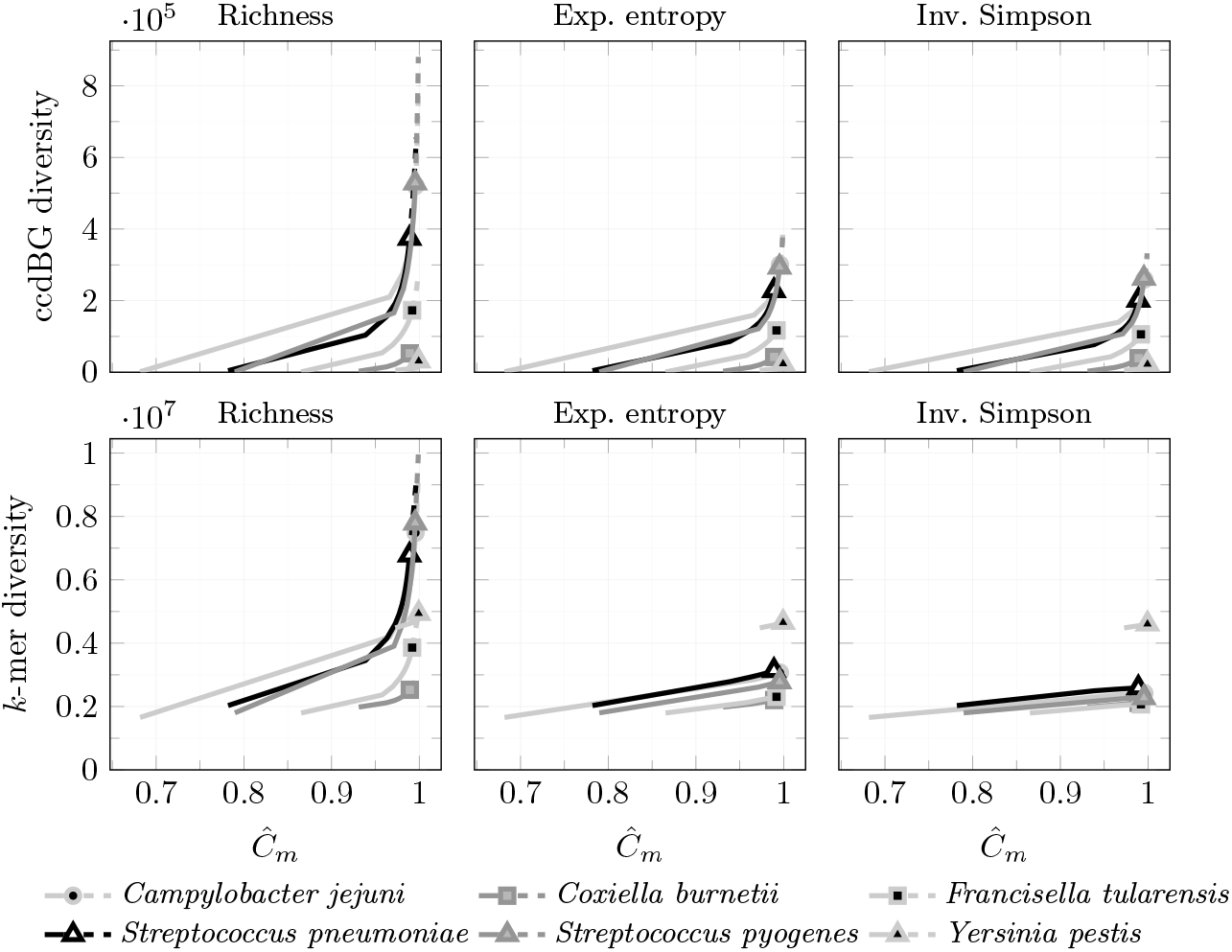
Hill numbers of ccdBG and *k*-mers for the group of species with at most 1 billion distinct *k*-mers over coverage.

Note that this division is based on the total number of distinct *k*-mers and not on the length of the genomes. For example, *C. jejuni* and *H. pylori* have similar genome lengths, but are in two different groups.

In the first group, *C. botulinum* and *H. pylori* show very similar graph diversity, despite *C. botulinum* having more than twice the genome size of *H. pylori* and seven times more *k*-mer richness. Their graphs are similar both in the number of unitigs and how evenly they are colored.

*B. aphidicola, P. marinus*, and *R. palustris* appear more diverse for their coverage, but strict conclusions are difficult due to their relatively low coverage, ranging from 34% to 67%.

In the second group, all species show higher coverage, which aligns with their lower observed diversity. The ranking is consistent between graphs and *k*-mers, with the exception of *Y. pestis*. Although coverage is intended to improve comparisons between species with different genome sizes, the relatively large genome of *Y. pestis* (about 4.7 Mbp) compared with the other species in this group (around 2 Mbp) inflates its *k*-mer diversity. In contrast, the ccdBG places it as the least diverse species, which is consistent with the fact that *Y. pestis* has limited variation and is largely clonal (Girard et al. 2004).

### 7.1 Pangenome openness

Finally, we characterize the growth of the 12 bacterial pangenomes using Heaps’ law. Following Tettelin et al. (2005), we fit the expected number of new unitigs added at genome *m* with the model *Km*^−*α*^. The exponent *α* is commonly used to quantify pangenome *openness*: smaller values indicate that new elements continue to be discovered at a higher rate as more genomes are added.

For ten species the adjusted *R*^2^ of the fit was between 0.954 and 0.997. *R. palustris* has a weaker fit (*R*^2^ = 0.608), likely due to its small sample size. *B. aphidicola* and *P. marinus* show an increasing number of new unitigs with *m*. This pattern is consistent with new *k*-mers breaking existing unitigs faster than new unitigs are formed at these sample sizes. Their fitted *α* is therefore negative, and we exclude them from the comparison of openness exponents.

Figure 7 compares the fitted exponents *α* from *k*-mer and ccdBG richness for the nine remaining species. The two sets of estimates follow a similar trend, but ccdBG *α* is lower than the *k*-mer *α* for most species, suggesting that graph compaction reduces the apparent degree of openness. Individual growth curves and fit diagnostics are reported in Appendix D.2.

**Figure 7.**
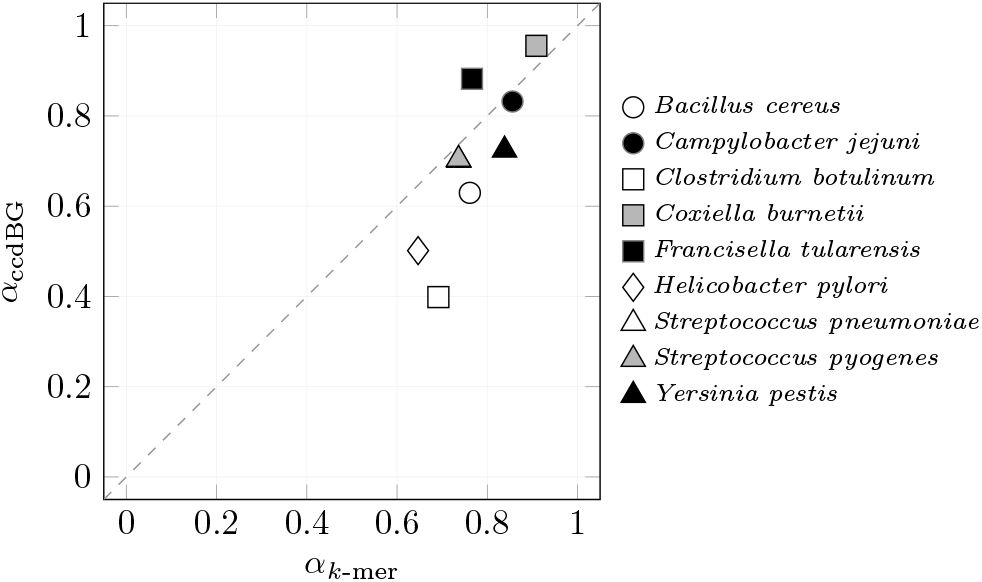
Comparison of Heaps-law exponents *α* estimated from *k*-mer richness (*x*-axis) and ccdBG richness (*y*-axis) for nine bacterial species. The dashed line marks *α*_*k*-mer_ = *α*_ccdBG_.

**Figure 8.**
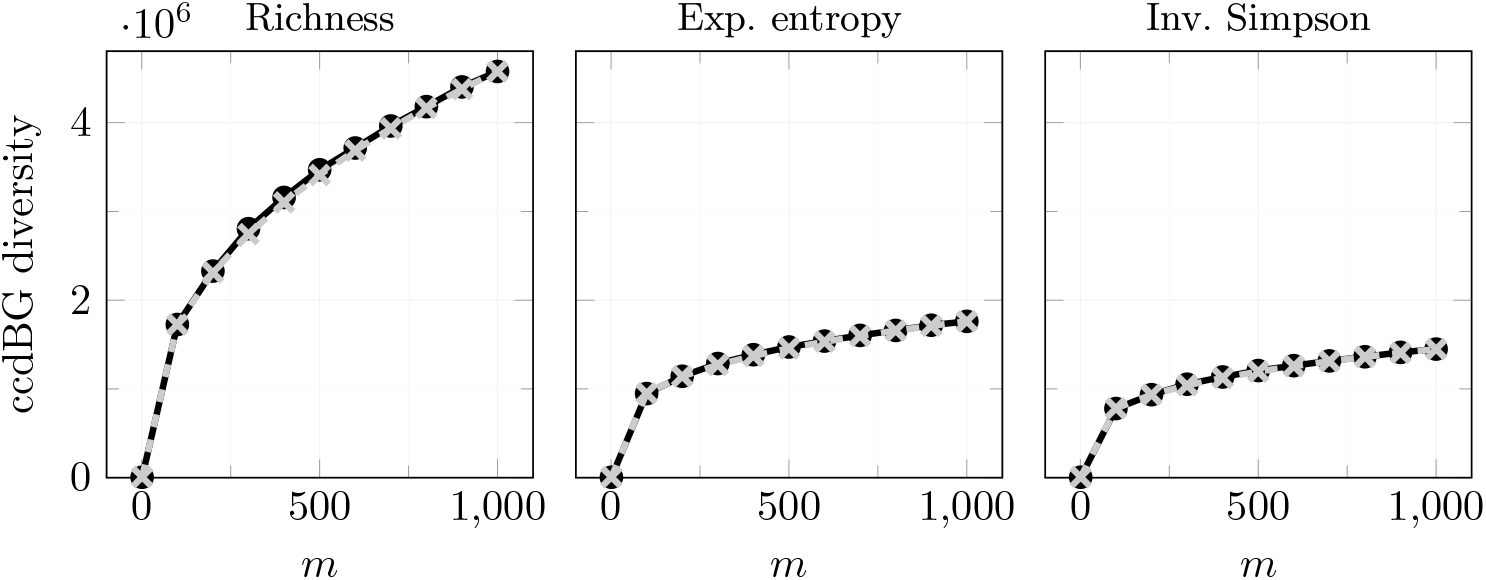
Interpolated Hill numbers for 1000 *E. coli* genomes. Solid black dots show the Hill numbers computed with the average unitig histogram over 10 repeated runs using Bifrost. The gray crosses show the sequence-based hypergeometric estimates.

**Figure 9.**
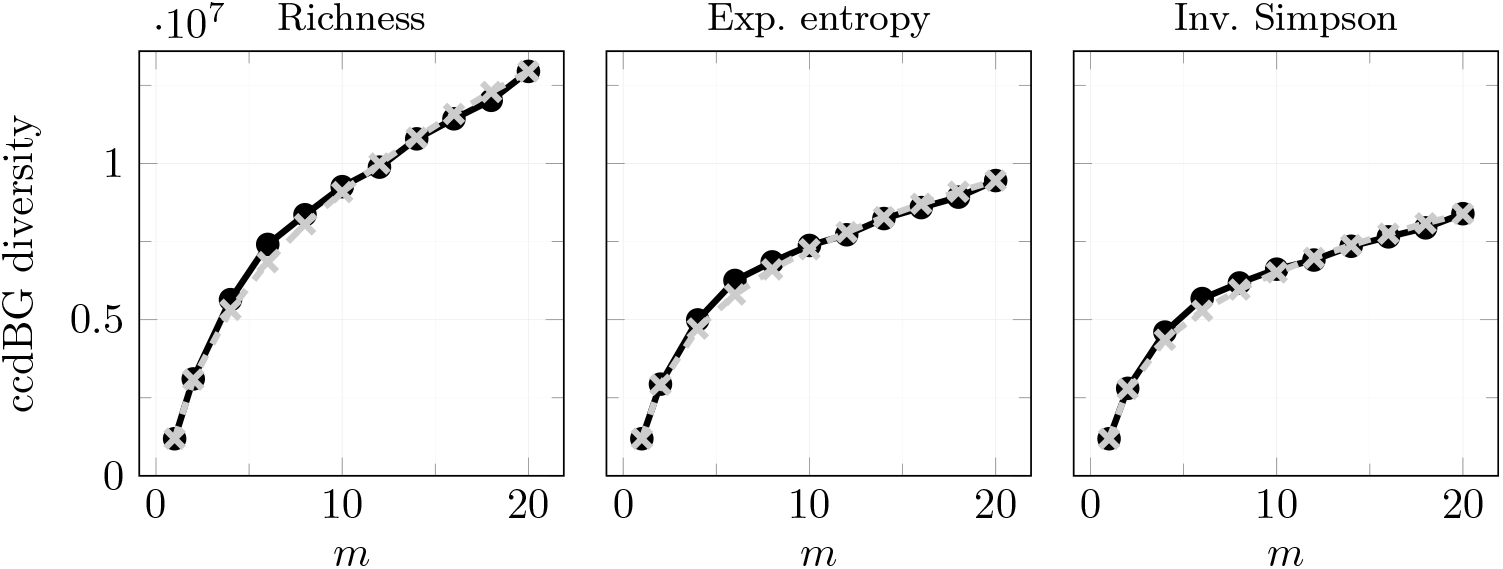
Interpolated Hill numbers for 20 *A. thaliana* genomes. Solid black dots show the Hill numbers computed with the average unitig histogram over 10 repeated runs using Bifrost. The gray crosses show the sequence-based hypergeometric estimates.

**Figure 10.**
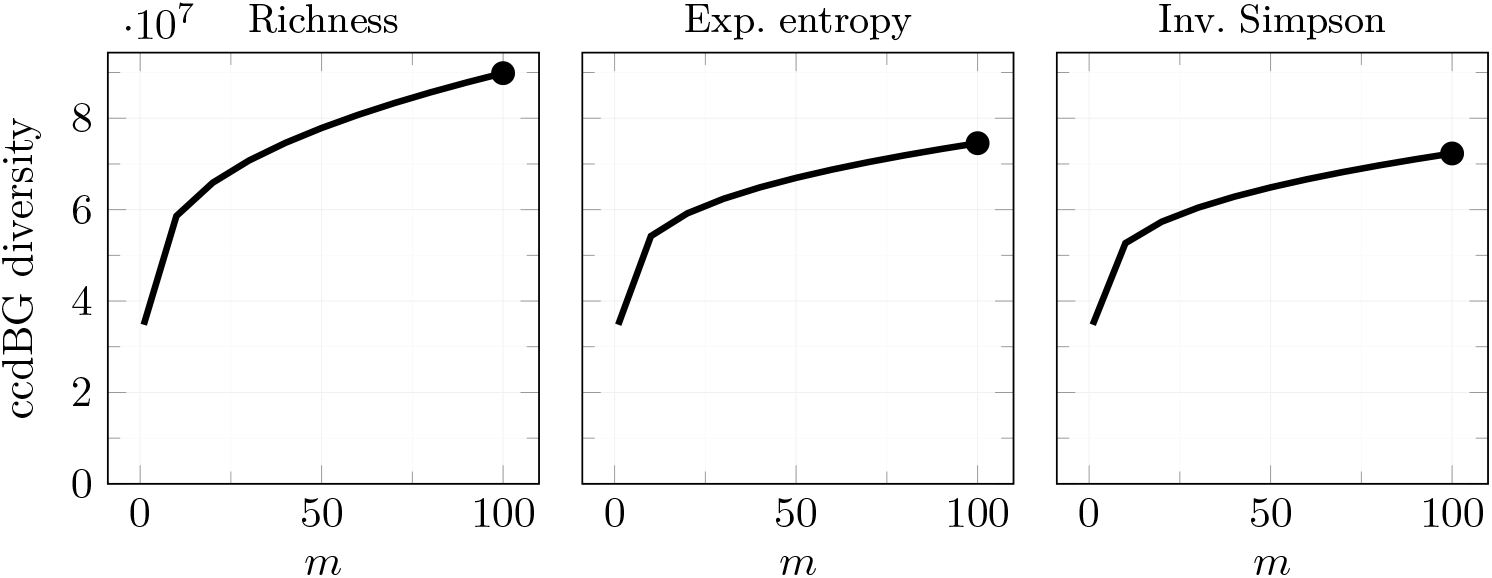
Hypergeometric interpolation of graph-based Hill numbers for the human pangenome.

**Figure 11.**
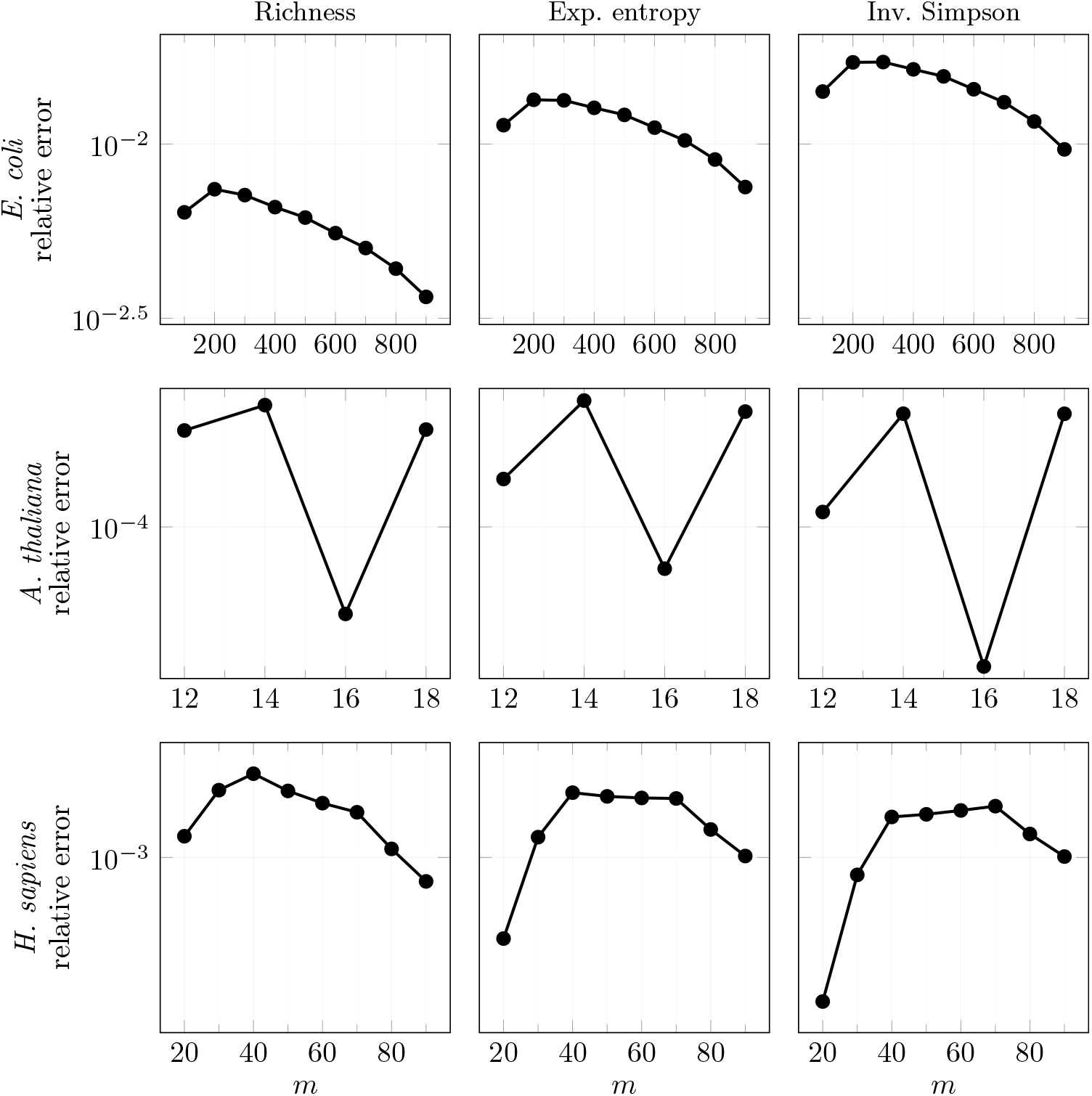
Relative error of adaptive Bernoulli-hybrid interpolation with *R* = 20 and *τ* = 0.001 against hypergeometric interpolation for 1000 *E. coli* genomes, 20 *A. thaliana* genomes, and 100 human genomes (top to bottom). Columns show richness, exponential entropy, and inverse Simpson index.

## 8 Discussion

The construction and analysis of graphs representing pangenomes is an emerging research direction. These graphs encode not only sequence information but also the distribution of genomes across the graph structure. However, they are rarely built from the same number of genomes, and even when they are, differences in genome size and underlying diversity can significantly affect diversity estimates. This makes direct comparison between pangenome graphs challenging and potentially misleading.

To make such comparisons meaningful, it is essential to normalize for these differences. Here, we introduced a novel method to interpolate and extrapolate Hill numbers over colored compacted de Bruijn graphs, a well-established class of graphs for pangenomes. These graphs are clearly defined, widely used, and supported by extensive research.

We showed that our approach can accurately estimate expected values of Hill numbers in polynomial time, avoiding the need for repeated graph constructions. We also applied this method to compare the ccdBG-based Hill numbers of the 12 bacterial species, for matching levels of *k*-mer coverage.

Future work could extend this framework to metagenomic data, for example by treating MAGs or metagenomic reads as colors, but requires careful definition of the sampling unit and construction of colored compacted de Bruijn graphs at metagenomic scale.

## 9 Acknowledgments

This project received funding from the European Union’s Horizon 2020 research and innovation programme under the Marie Sklodowska-Curie grant agreement No 956229. It was also supported by the BMBF-funded de.NBI Cloud within the German Network for Bioinformatics Infrastructure (de.NBI) (031A532B, 031A533A, 031A533B, 031A534A, 031A535A, 031A537A, 031A537B, 031A537C, 031A537D, 031A538A), and the French Inria Challenge “OmicFinder”.

We thank Cinzia Pizzi and Jens Stoye for reviewing the thesis associated with this work and for their valuable feedback.

## 11 Data and Software Availability

Pangrowth is openly available at https://github.com/gi-bielefeld/pangrowth. Scripts, accession lists, and result files for the *E. coli, A. thaliana*, and human experiments are openly available at https://github.com/lucaparmigiani/pangenome-graph-diversity. The datasets for the 12-species bacterial comparison were previously described by Parmigiani et al. (2024).

## A Algorithm

To compute the interpolation ^*q*^Δ(*m*),1 ≤ *m* ≤ *N*, we need the frequencies *h*^*k*-mer^, *h*^uni-mer^ and *h*^infix^ of the pangenome.

The *k*-mer frequency *h*^*k*-mer^(*i*) can be found using a *k*-mer counter and then counting how many *k*-mers appear *i* times like described in (Parmigiani et al. 2024).

To compute the infix equivalents, which also contain the distribution of uni-mers, we process one genome at a time, extracting all (*k* + 1)-mers. Each (*k* + 1)-mer *x* is inserted into a hash table *H*, indexed by its infix *x*[2:*k*], to group together all infix equivalent (*k* + 1)-mers.

Each entry *H*[*x*[2:*k*]] is a tuple (*count, edge*_last_, *genome*_last_, *σ*). Each (*k* + 1)-mer *x* = *a x*[2:*k*] *b*, with *a, b* ∈ Σ, added to *H*[*x*[2:*k*]] has the same infix, so we only need to record the flanking characters *a* and *b*. The map *count*[*ab*] counts in how many genomes *x* occurs as a uni-mer. The value *edge*_last_ = *ab* corresponds to the last (*k* + 1)-mer seen, and *genome*_last_ is the genome from which it was last observed. These two fields are used to avoid counting the same edge multiple times within the same genome and finding when there are at least two infix equivalent (*k* + 1)-mers.

The value *σ* records the number of distinct genomes that contain at least one (*k* + 1)-mer with infix *x*[2:*k*]. If a genome contains at least two infix equivalent (*k* + 1)-mers, it contributes only once to *σ*. Algorithm 1 shows the pseudocode to create the hash table *H*. There are at most 16 infix equivalent (*k* + 1)-mers (4 possible values for *a* times 4 for *b*), plus 8 additional cases to represent the telomeres of the form *a*$ and $*b*. The addition of telomeres is performed at the beginning and at the end of the loop.

### Algorithm 1

Extracting (*k* + 1)-mers from genomes

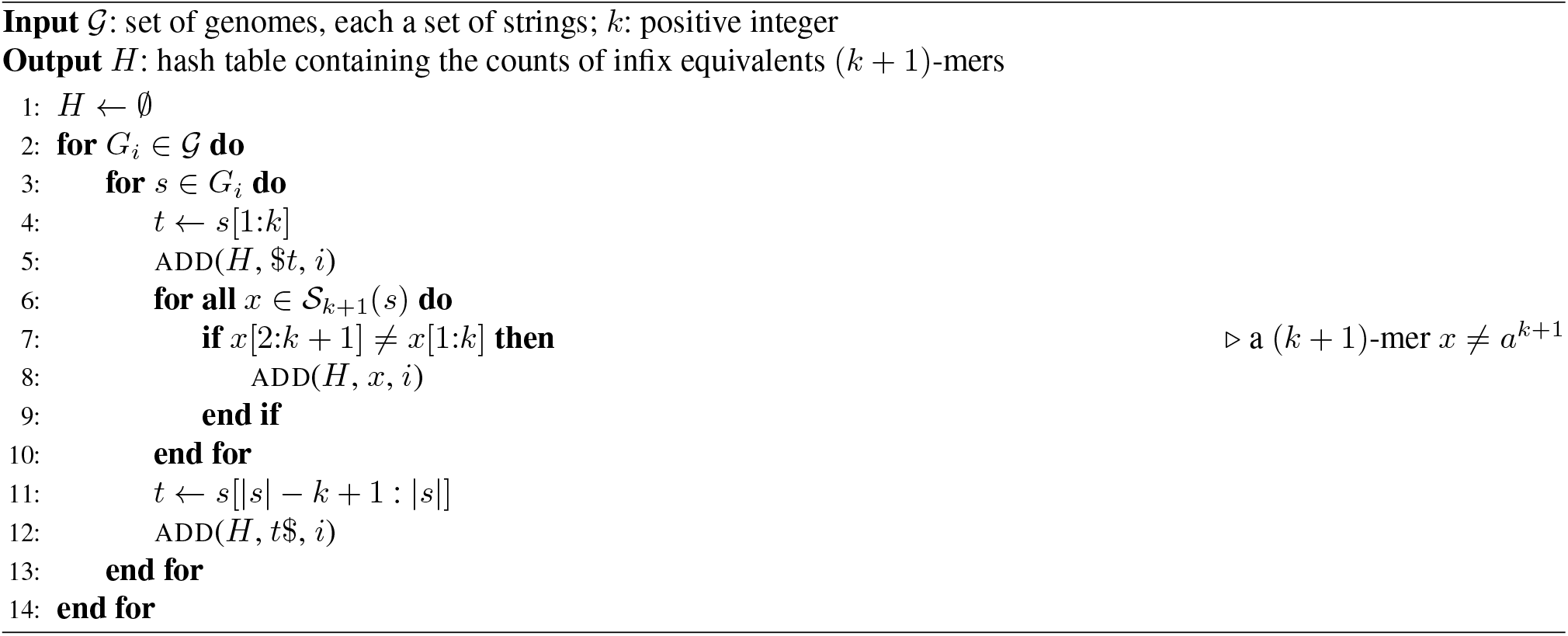

The procedure for adding *x* to *H* from a genome, denoted as ADD(*H, x, genome*), is illustrated in Algorithm 2. We first describe the case where *x* is a regular non-telomere (*k* + 1)-mer. After, we explain the insertion of a telomere.

The procedure begins by checking whether a record *r* for the infix *x*[2:*k*] already exists in *H*. If not, the record is created, and both the *count*[*ab*] (for the corresponding flanking characters) and *σ* are set to 1. When the record already exists, there are two main cases to distinguish: whether the current *genome* is the same as *genome*_last_, or not.

If the genome is different, *count* and *σ* are incremented. If the genome is the same and the new flanking edge differs from the previously stored one, it means that two infix-equivalent (*k* + 1)-mers were found in the same genome. In this case, the position can no longer form a uni-mer, so the *count* is decremented (lines 17–18), and *edge*_last_ is set to 1 to indicate it.

When adding a telomere, we do not increment *σ* immediately, since such a fragment alone cannot form a uni-mer. However, if a corresponding opposite telomere from the same genome is later found, the two can be joined to form a uni-mer. This is handled in lines 19–25 using the function UNITE(*edge*_last_, *edge*), which combines the partial edges. The function CONTAINED(*edge*_last_, *edge*) checks whether two (*k* + 1)-mers are compatible: either identical, opposite telomeres, or one being a prefix or a suffix of the other. Note that a telomere can break a (*k* + 1)-mer if it is not contained and the other way around.

Lastly, we create *h*^infix^ from *H*, as shown in Algorithm 3. For each infix equivalent (*k* + 1)-mer, we count the number of genomes that form a uni-mer if selected in the same subset *m* without the genomes that cannot be taken, represented by *σ*. The algorithm counts the frequency of these infix equivalents in lines 12–20. Lines 4–11 handle the special case of telomeres.

### Algorithm 2

Add a (*k* + 1)-mer *x* to the hash table *H*

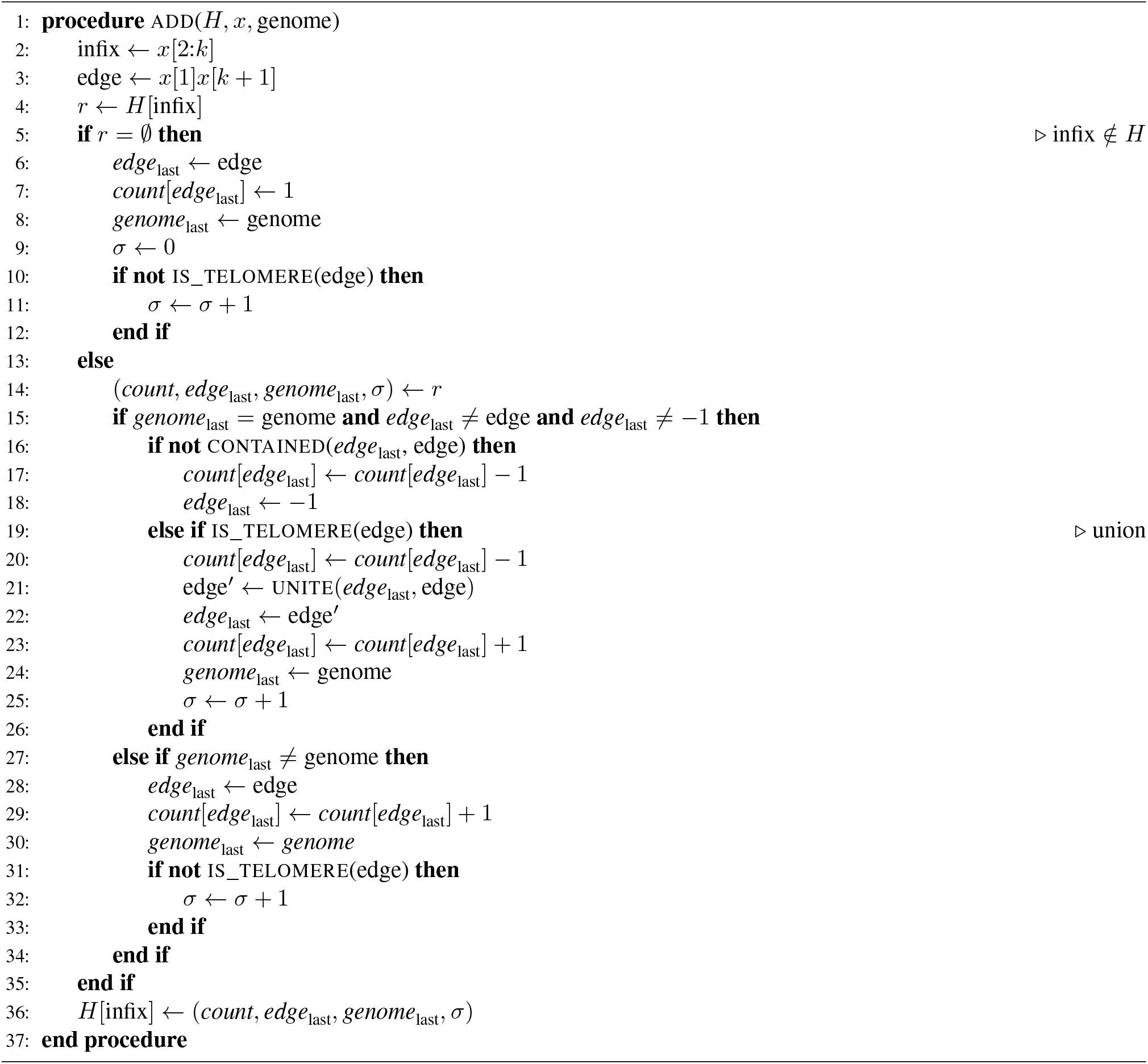

For example, if some genomes form a uni-mer represented by the flanking pair *ab*, then genomes containing a conflicting pair *a*^*′*^*b*^*′*^ (with *a* ≠ *a*^*′*^ or *b* ≠ *b*^*′*^) cannot be included in the same subset. However, genomes containing telomeres such as *a*$ or $*b* do not cause conflicts and can be included.

Computing ^*q*^Δ(*m*) for all possible *m*, 1 ≤ *m* ≤ *N*, with our approach requires *O*(*N* ^4^). For each *m* we need 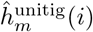. To compute it we need the expected *k*-mer frequencies, which take *O*(*m*^2^) for each value of 1 ≤ *m* ≤ *N*, so in total *O*(*N* ^3^), and the expected uni-mer frequency which takes *O*(*m*^3^), so *O*(*N* ^4^). In practice it is common to compute ^*q*^Δ(*m*) only for a subset *s* ∈ [1, *N*] of points, usually *s* = 30 (Hsieh et al. 2016), bringing down the complexity to *O*(*sN* ^3^).

### Algorithm 3

Populate histogram *h*^infix^ from hash table *H*

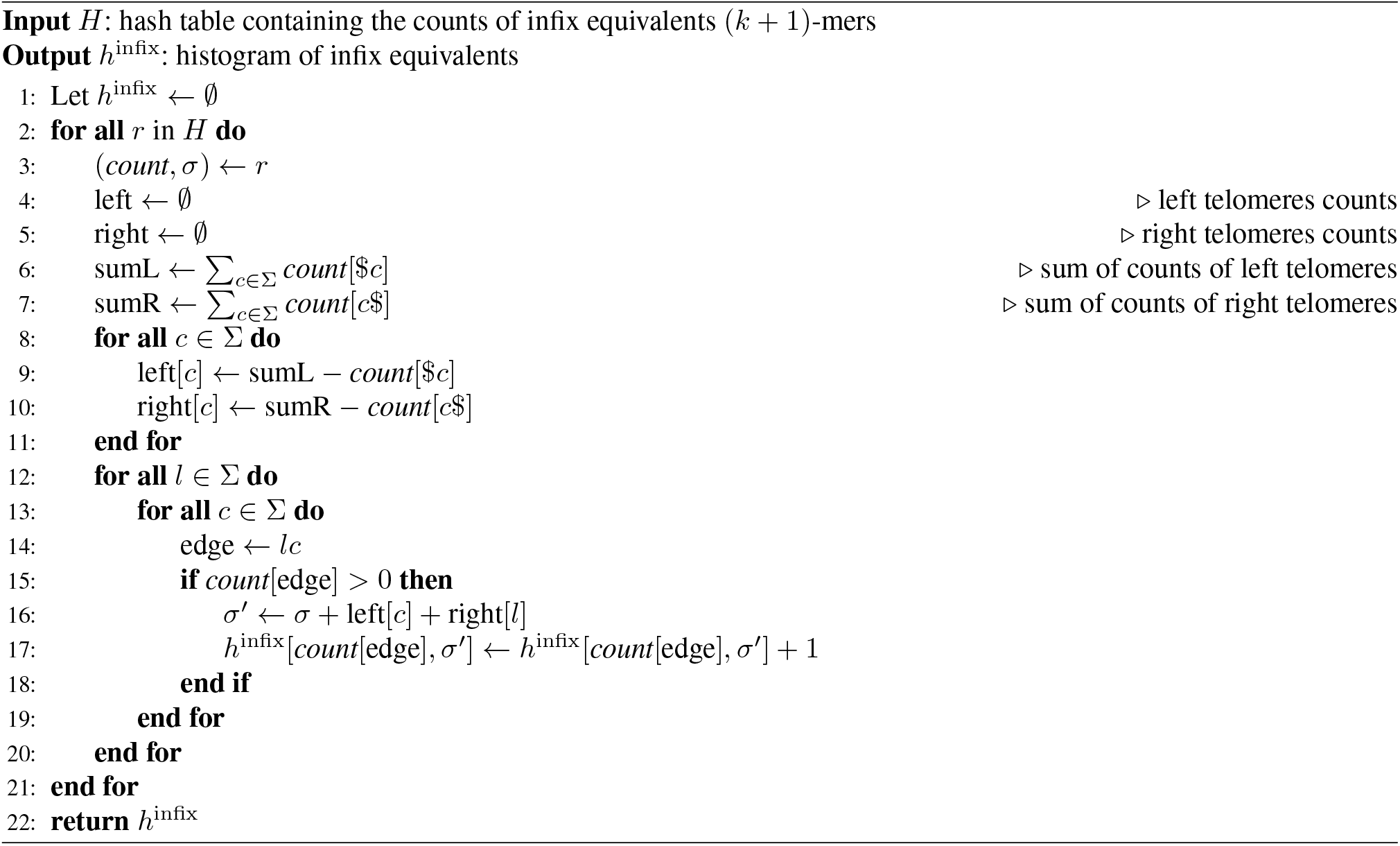

### A.1 Canonicity

We now extend these ideas to the canonical ccdBG. The key concept is to define an orientation of a (*k* + 1)-mer based on its infix. To ensure a consistent orientation we always process a (*k* + 1)-mer in the direction where the infix is lexicographically smaller than its reverse complement, so we consider *x*, if *x*[2:*k*] ≤_lex_ *x*[2:*k*]^*rc*^ or *x*^*rc*^, otherwise.

All the algorithms described previously remain unchanged, except for Algorithm 1, which must now account for directionality. The modified version is shown in Algorithm 4.

When inserting a telomere *t* of the form $*t* (lines 4–9), we check whether the suffix *t*[1:*k* − 1] is lexicographically smaller or equal than its reverse complement. If it is, we keep it as $*t*; otherwise, we insert *t*^rc^$. A similar logic is applied when handling telomeres of the form *t*$ (lines 19–24).

The second difference with Algorithm 1 is in the insertion of the (*k* + 1)-mer, not only do we need to take the canonical direction but also avoid adding any (*k* + 1)-mer *x* such that their infix is the same as its reverse complement, since these instances will not form a uni-mer.

#### Algorithm 4

Extracting *canonical* (*k* + 1)-mers from genomes

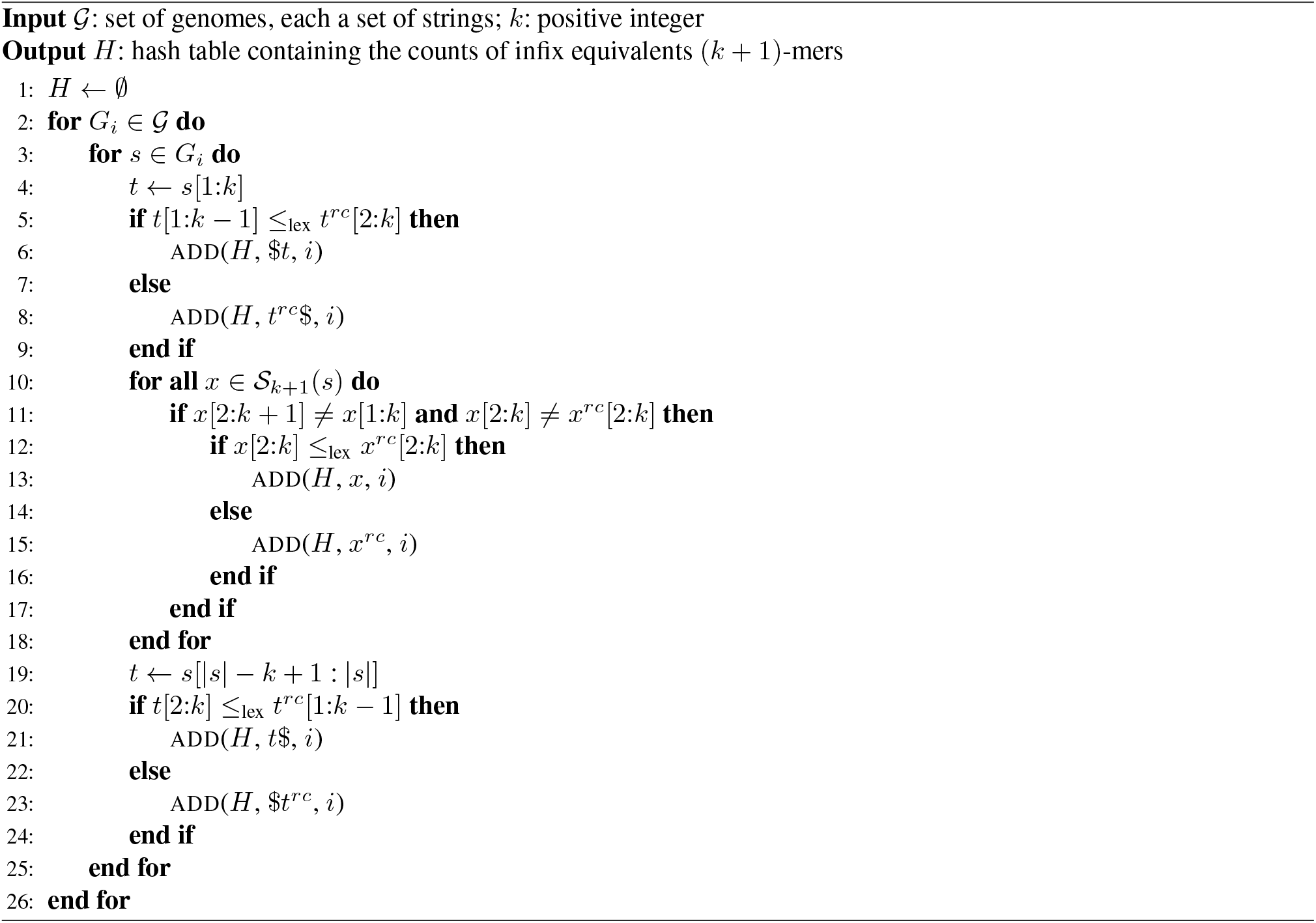

## B Interpolation

For Bifrost and GGCAT, we built the ccdBG for different subsamples of the data. This was repeated 10 times using different subsamples, except for *m* = *N* where only one sampling is possible. Then, we took the average from the resulting unitig histograms to compute Hill numbers.

Bifrost supports incrementally updating an existing graph using the update command. To ensure a fair comparison, we used Bifrost API to avoid writing each newly constructed graph to disk, as only the histograms were needed. We started from a graph built on a single genome and incrementally added more genomes in batches. Bifrost’s default behavior is to build a new graph from the new genomes and merge it with the existing one, therefore we kept the same functionality.

GGCAT was run from scratch each time, for each subsample. Since a core feature for GGCAT is to use disk storage to reduce RAM usage, we kept this default behavior.

For each dataset, we sampled Hill numbers at 11 points. For *E. coli*, we started from a single genome, and then from 100 to 1000 in steps of 100. For *A. thaliana*, we started from one and then from 2 to 20 in steps of 2.

### B.1 Human pangenome resources

### B.2 Bernoulli-hybrid interpolation

The maximum relative errors across interpolation points were 0.742%, 1.339%, and 1.719% for *E. coli*; 0.063%, 0.068%, and 0.056% for *A. thaliana*; and 0.163%, 0.146%, and 0.135% for the human pangenome, for richness, exponential entropy, and inverse Simpson index, respectively.

## C Uni-mers Extrapolation

We now describe the two theoretical processes that determine the expected number of uni-mers. Let *Z*(*x*) be the set of (*k* + 1)-mers that are infix equivalent to *x*, namely *Z*(*x*) = {*z*|*z* ∈ ℰ_*k*+1_(*G*) ∧ *z*[2, *k*] = *x*[2, ∧ *k*] *z* ≠ *x*}. Analogous to Equation (1), we can write a theoretical expression for uni-mer growth. Here, is the set of all possible (*k* + 1)-mers in the pangenome, even the ones not yet seen. Then:

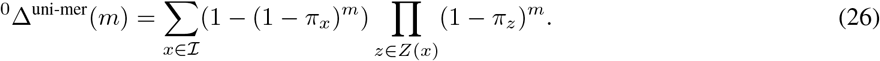

This represents the number of (*k* + 1)-mers that appear in at least one of the *m* genomes and have no other infix equivalent. Note that this holds for any positive integer *m*, not only the one restricted to 1 ≤ *m* ≤ *N* . From Equation (26) we compute the difference ^0^Δ^uni-mer^(*m* + 1) − ^0^Δ^uni-mer^(*m*) as

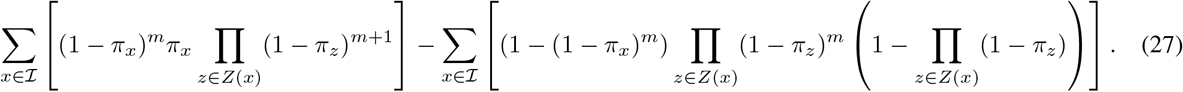

### C.1 Random-holdout evaluation

For each observed fraction, every subset was extrapolated to the complete dataset size and compared with the Hill numbers computed from all genomes. Figure 12 reports the signed relative error across the 10 randomized subsets.

**Figure 12.**
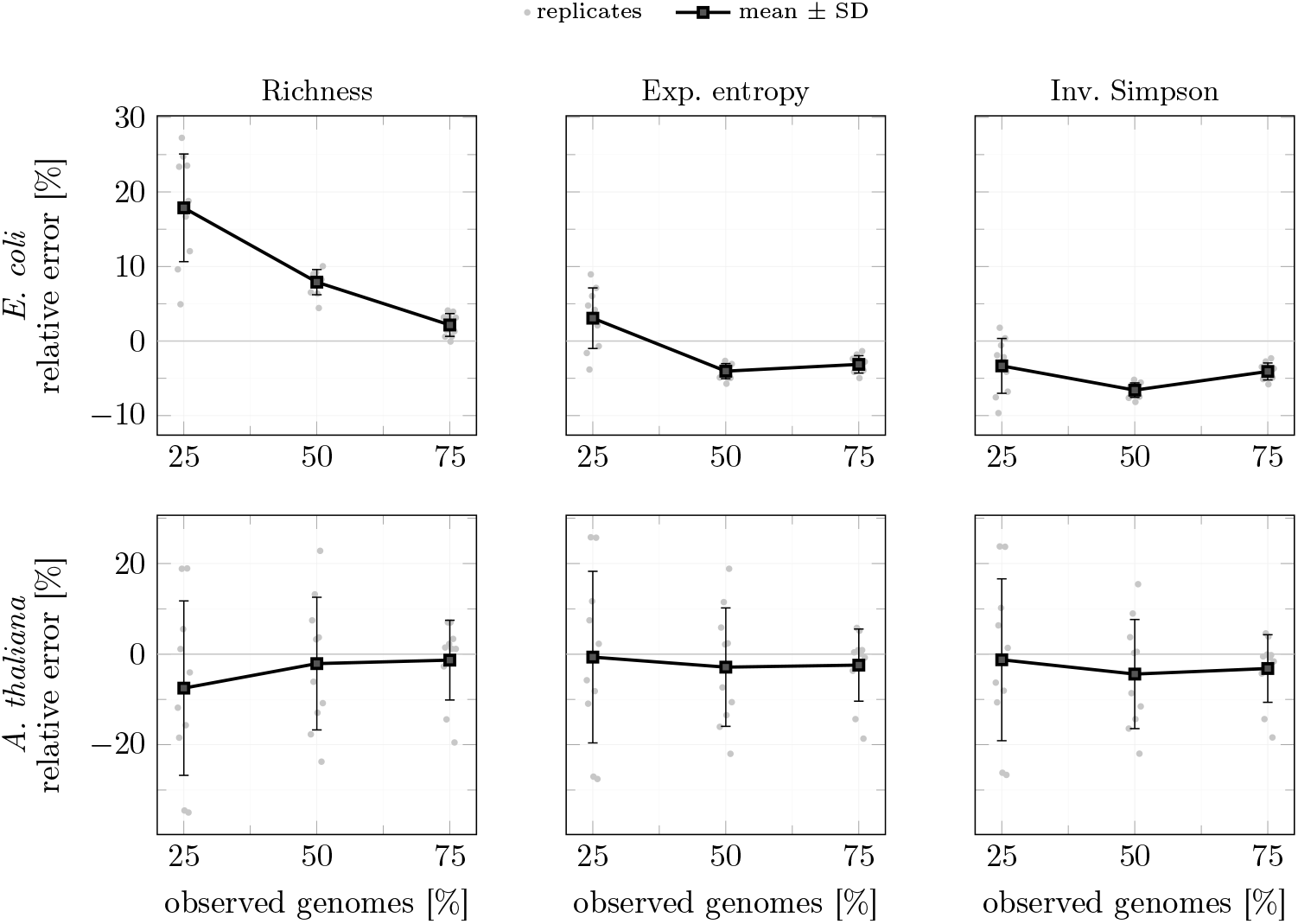
Signed relative error of the extrapolated Hill numbers at the complete dataset size for 1000 *E. coli* genomes (top) and 20 *A. thaliana* genomes (bottom). Gray points represent individual random subsets; black squares and error bars show the mean and one standard deviation. Columns show richness, exponential entropy, and inverse Simpson index.

**Figure 13.**
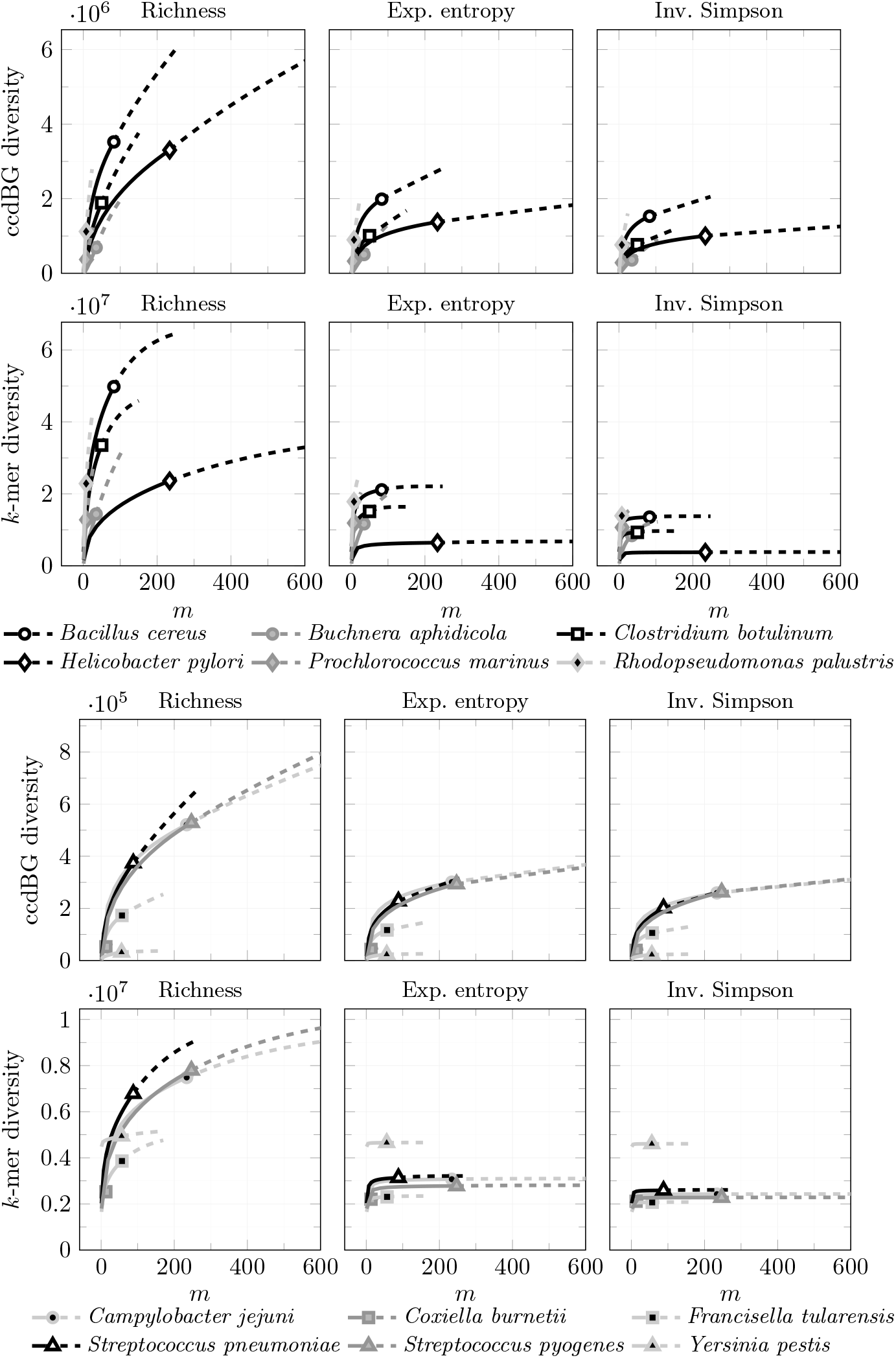
Hill numbers of ccdBG and *k*-mer for the group of species with more than 1 billion distinct *k*-mers (above) and with at most 1 billion (below).

**Figure 14.**
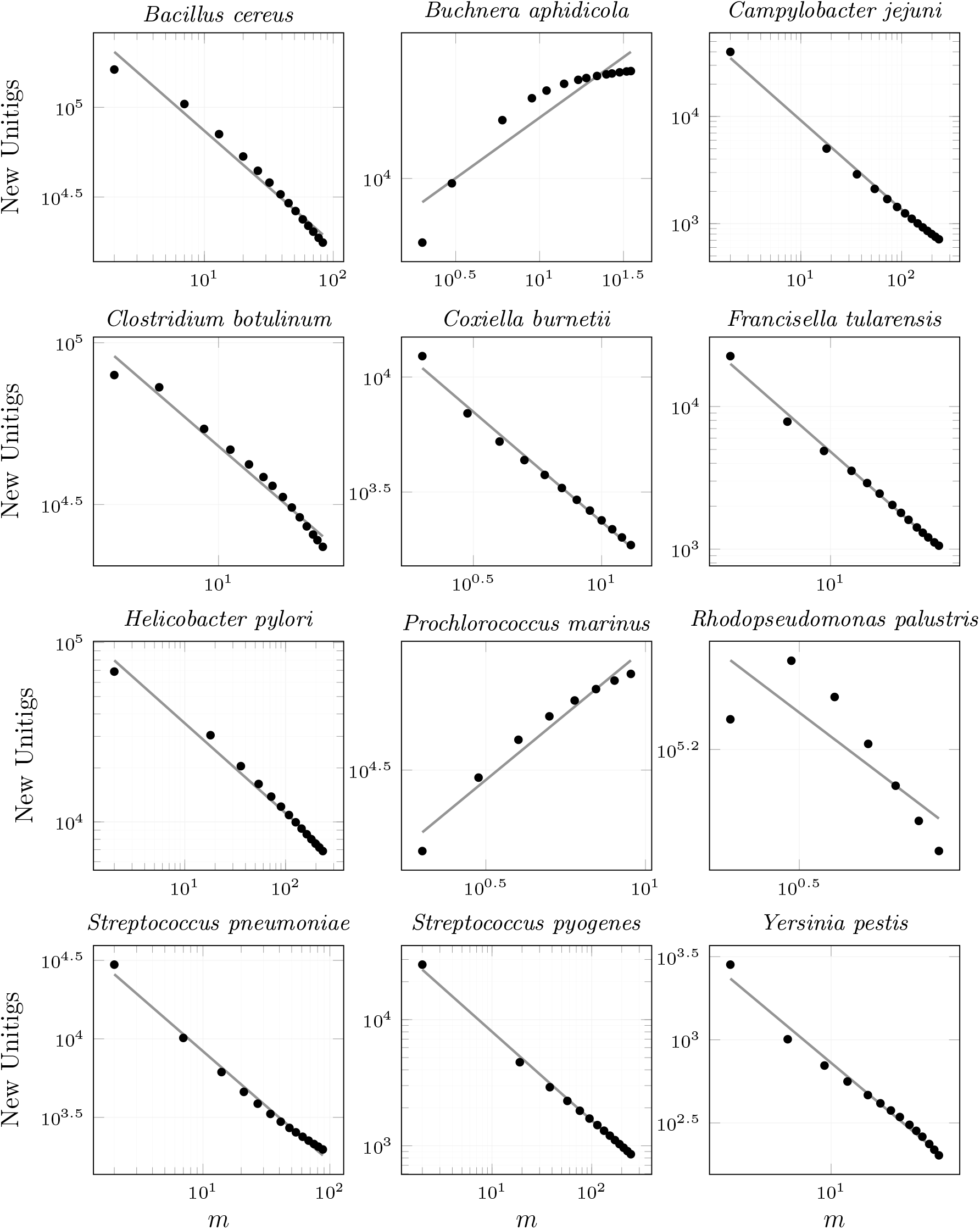
Heaps-law fits for ccdBG unitig richness for each of the 12 bacterial species. Gray lines show the fitted model *Km*^−*α*^; black dots show the observed new-unitig counts at the selected genome counts.

## D Pangenome Graph Comparison

In both groups, and for both ccdBG and *k*-mer, richness appears to grow indefinitely, with the exception of *Y. pestis*, where it levels off quickly. For the exponential entropy and the inverse Simpson index, the *k*-mer reaches an asymptotic behavior in all cases. This indicates that growth is primarily driven by unique *k*-mers, while those shared across multiple genomes are mostly already present.

In the second group (species with at most 1 billion *k*-mers), the exponential entropy and the inverse Simpson index values are very close, suggesting a more even distribution of *k*-mers and unitigs. In contrast, the first group shows a noticeable drop from entropy to inverse Simpson index, indicating a less even distribution.

While ccdBG and *k*-mer curves generally show similar trends, they are not identical. In the first group, the relative order of species remains the same for both, except for *H. pylori*. Here, the *k*-mer Hill numbers rank lower than the ccdBG values (below *P. marinus*), while the ccdBG ranks higher. This suggests more structural variation in the graph than what is reflected by the *k*-mer count alone.

Similarly, in the second group, the relative species order is consistent except for *Y. pestis*. It ranks fourth in richness and first in both exponential entropy and inverse Simpson index for *k*-mers, but drops to last place when measured on the ccdBG. The higher richness in *k*-mers is due to its relatively large genome size compared to the other species. However, its pangenome grows slowly. This highlights the value of using pangenome graphs: the compaction of *k*-mers acts as a normalization step that reveals differences not captured by *k*-mer counts alone.

### D.1 Unitig length distributions

### D.2 Pangenome openness

## E Coverage

We define the coverage for *k*-mers to compare two ccdBGs with the same *k*-mer coverage. Due to the *k*-mer distribution (Equation (1)) we can use the estimators defined by (Chao and Jost 2012). For *N* genomes the estimated coverage is:

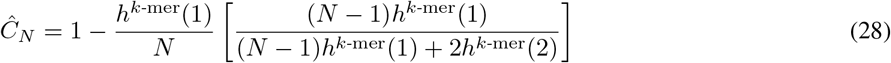

The interpolated coverage for *m < N* is:

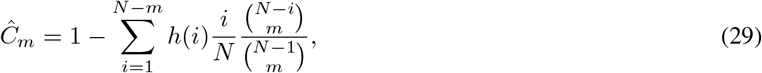

and the extrapolated coverage for *N* + *m*^∗^ is:

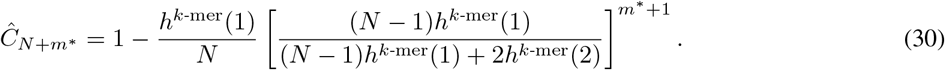

